# An integrative modelling approach to the mitochondrial cristae

**DOI:** 10.1101/2024.09.23.613389

**Authors:** Chelsea M. Brown, Marieke S. S. Westendorp, Rubi Zarmiento-Garcia, Jan A. Stevens, Sarah L. Rouse, Siewert J. Marrink, Tsjerk A. Wassenaar

## Abstract

Mitochondria are implicated in many cellular functions such as energy production and apoptosis but also disease pathogenesis. To effectively perform these roles, the mitochondrial inner membrane has invaginations known as cristae that dramatically increase the surface area. This works to provide more space for membrane proteins that are essential to the roles of mitochondria. While separate components of this have been studied computationally, it remains a challenge to combine elements into an overall model. Here we present a workflow to create a comprehensive model of a crista junction from a human mitochondrion. Our coarse-grained representation of a crista shows how various experimentally determined features of organelles can be combined with molecular modelling to give insights into the interactions and dynamics of complicated biological systems. This work is presented as an initial ‘living’ model for this system, intended to be built upon and improved as our understanding, methodology and resources develop.

## Introduction

Mitochondria, colloquially known as the powerhouse of the cell, are organelles that are found in almost all eukaryotic cells and are vital not only for energy generation, but also for cell signalling, metabolism and apoptosis^1,2^. It has an unusual morphology for eukaryotic organelles, consisting of two membranes: the inner (IMM) and outer (OMM) mitochondrial membranes. The OMM resembles a typical outer bilayer for a cell, but the IMM has a complex topology comprising of large invaginations of the membrane called cristae^3^. The shape of the IMM serves to increase the surface area of the membrane, providing more space for the critical proteins it houses such as the respiratory complexes and ATP synthases^4^, which are vital for the metabolic functions that mitochondria perform.

The shape of the membrane is closely regulated by various proteins, such as the mitochondrial contact site and cristae organising system (MICOS), ATP synthases (ATPase) and the long isoform of Optic atrophy protein 1 (l-OPA1)^5,6^. The lipid composition also reflects the curved nature of the IMM, comprising a high level of cardiolipin (CDL2) which can also aid in maintenance of the membrane shape^7,8^.

Due to the relevance to human disease^1,9^, mitochondria have been extensively studied experimentally^10^. This has resulted in many protein structures^11–19^, sub-cellular location analysis^20,21^, and databases quantifying absolute amounts of mitochondrial proteins^22^. While these studies provide a vast amount of information about mitochondria and their components, the dynamics and molecular interplay between them are still poorly understood on a cellular level.

Molecular dynamics (MD) has proved itself as an extremely powerful tool to get molecular level insight into crucial biological processes and has been referred to as ‘Computational microscopy’^23,24^. Computational studies of specific mitochondrial proteins and lipids are not new^8,18,25–30^ with many of the most important complexes being widely reported. But often these simulations are done in isolation from other proteins, lipid curvature or the presence of a dual lipid bilayer system. Larger scale studies have been applied to the mitochondria such as ATPase dimer effects on membrane curvature^6^, cardiolipin binding to respiratory chain supercomplexes^28^ and a lipid only model of a mitochondrion^31^.

However, as computational power continues to increase, simulations of highly complex systems are now being realised. This brings rise to ‘*in-situ*’ modelling^32^ with high levels of detail, such as the JCVI-Syn3A minimal cell^33^ and virus particles^34,35^.

To make the most accurate representation of a biological system, integrative modelling can be employed^32,33,35^. This aims to bring together a vast collection of experimental and computational data to create a reflective model. The constituent elements will be fed into a high-throughput pipeline to examine individual behaviour. When assembling large systems, it is increasingly important to start close to equilibrium behaviour (i.e. how it would exist biologically) as this will be difficult to achieve with systems of such complexity in timescales that are currently available.

Herein, we present a representative model of the human mitochondrial cristae, based on the latest version of the popular Martini coarse-grain (CG) force field^36^. The model results from integrating a range of data from experimental and theoretical studies and includes both membranes, curvatures that are found in the organelle and 113 unique protein chains resulting in 96,580 residues in the system. These structures represent accurate oligomerisation states and only using mature sequences. This is presented as an initial representation and is intended to be a ‘living model’ which can be updated as more experimental data emerges.

## Results

### Selection and preparation of protein structures

To decide which proteins to include in our model of a human mitochondrial crista, the first step was performing a vast literature search. We focused on those proteins/complexes with high abundance, with most identified as key to maintenance of cristae morphology, components of the respiratory chain or proteins for peptide translocation. This resulted in the selection of 13 proteins/complexes, listed in Table 1. The next step involved setting up suitable models of these proteins to be included in the simulation. We began by selecting suitable starting points for the structures, most often from the Protein Data Bank (PDB). When the structures of human proteins were available, these were completed to include any missing residues. This was cross-referenced with literature to ensure the right oligomerization state was represented. If the human protein structure had not been reported, we employed homology modelling^37^ or AlphaFold^38^ structure prediction. The method used for each can be seen in SI Table 1.

**Table 1:**
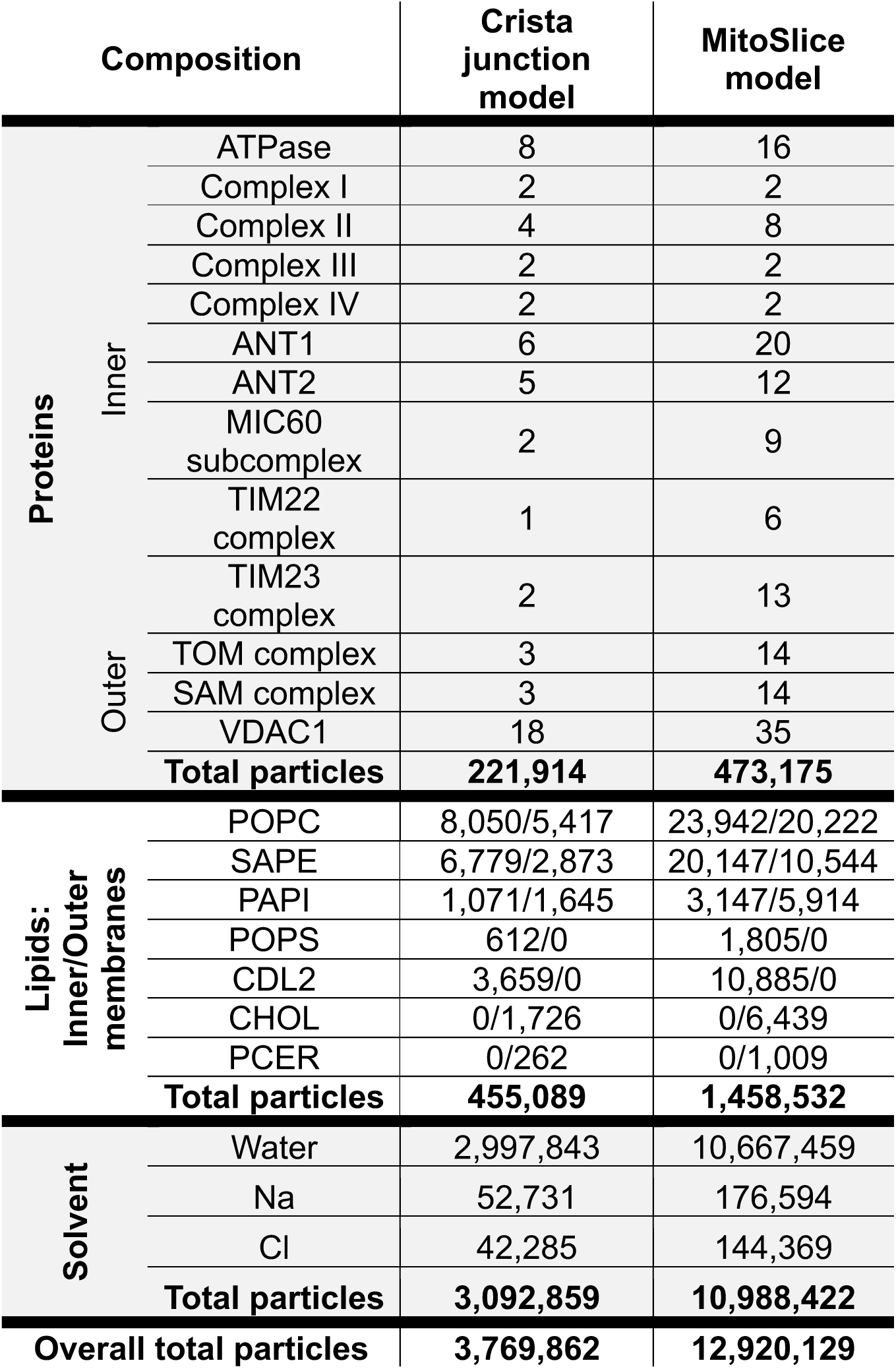
Compositions of the mitochondrial models produced in this study.

One of the most widely reported proteins to influence cristae shape is the ATPase^6,39^. When the complex exists as a monomer there is no significant influence on membrane shape, but when the dimer is formed curvature is induced that has been observed both experimentally^12,39^ and computationally^6^. A structure of a human ATP synthase complex has recently been solved^40^ (PDB ID: 8H9S), but only as a monomer. As the dimer is critical for maintenance of the cristae curvature, this is required modelling. A dimer has been solved for another mammalian species (Bovine, PDB ID: 7AJB)^17^ and we used this as a template to build the model for the human dimer (Figure 1A). This ensured that the angle between the dimers in this model reflected those seen in mammalian mitochondria.

**Figure 1:**
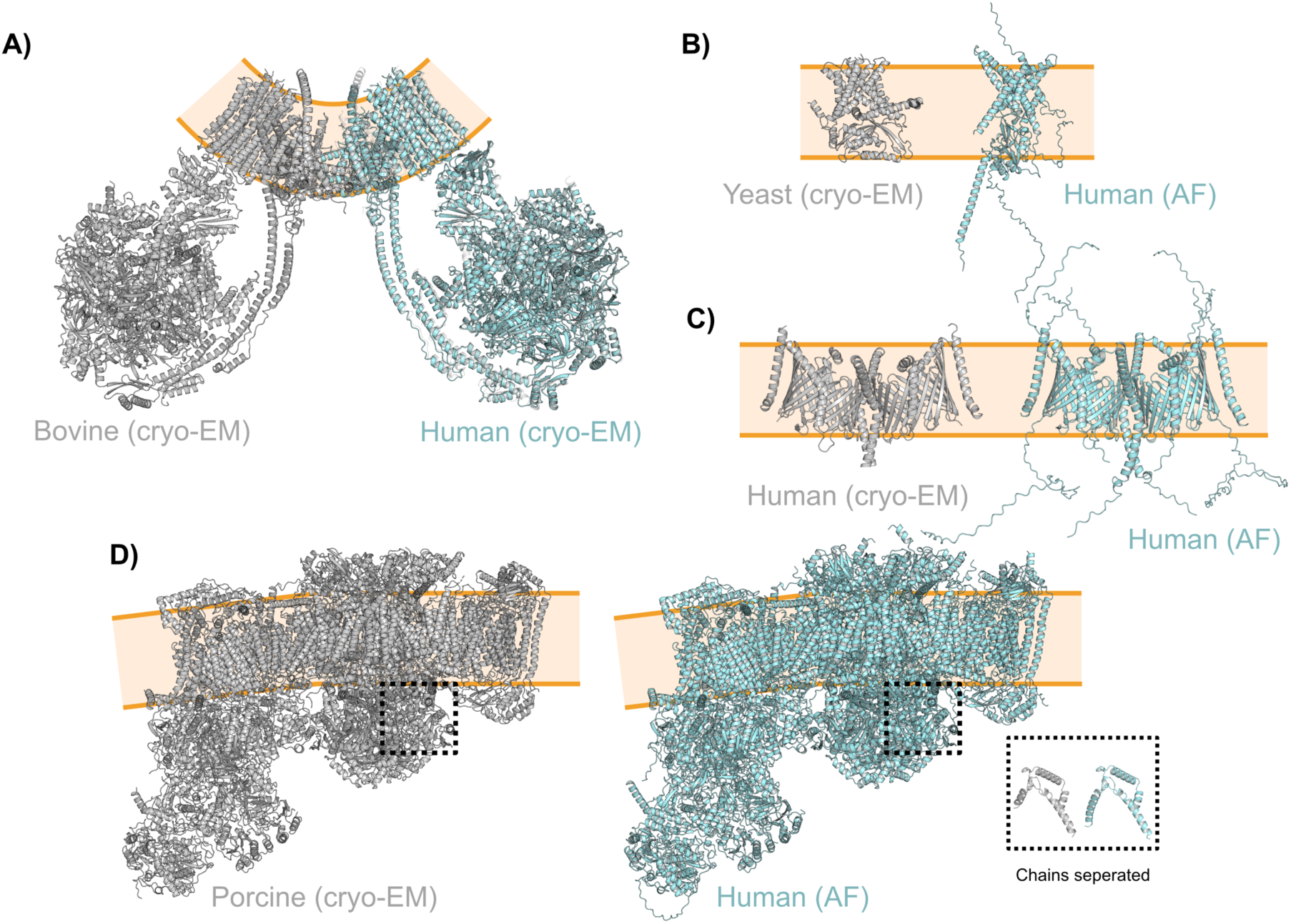
Protein structure generation examples. (A) The human ATPase structure (PDB ID: 8H9S, shown in blue) overlayed with the bovine ATPase structure (PDB ID: 7AJB, shown in grey). The position of the membrane is indicated with orange lines. (B) The AlphaFold (AF) model of the human TIM23 complex (shown in blue) with the solved yeast TIM23 complex (PDB ID: 8SCX, shown in grey). (C) The altered structure of the human TOM complex with added residues using AlphaFold (shown in blue) and the experimentally resolved structure (PDB ID: 7VD2, shown in grey). (D) The modelled human respiratory supercomplex using AlphaFold (shown in blue) with the solved porcine structure (PDB ID: 8UGH, shown in grey). An inset of one chain from each is shown for direct comparison.

Some of the most widely studied mitochondrial proteins are those of the respiratory chain. Complexes I, III and IV form a super complex^19,41^ while complex II is reported to exist separately^13,42^. Recently, a study showing the main oligomeric state of the super complex in mammalian mitochondria was published^11^, and ‘Type A’ of this porcine superstructure (PDB ID: 8UGH) was used as a template to construct a human homologue. Manual generation of the human structure was completed by identifying homologous chains using UniProt^43^, then downloading the pre-generated AlphaFold2^38^ predictions; removing residues labelled as signal peptides (as per UniProt^43^). We then aligned these structures with the relevant chain from the porcine structure. This was successful for all but 2 of the 81 chains, both of which belong to Cytochrome b-c1 complex subunit Rieske. When the subunits were then combined, the RMSD was measured to be 0.558 Å (60450 of 78235 atoms, after 5 cycles of refinement), the comparison can be seen in Figure 1D. The structure of the human respiratory complex II has been resolved^13^ (PDB ID: 8GS8) but not all residues are identified. To complete the structure, a workflow similar to that used for the super complex was employed. The resulting RMSD between the original structure and the modified version was 0.548 Å (4122 of 4901 atoms, after 5 cycles of refinement). The method used here ensures inclusion of the disordered terminal regions of each subunit (Figure 1D), which are not resolved experimentally.

To transport ATP/ADP through the inner membrane of the mitochondria, there are two isoforms of the main transporter: ADP/ATP translocase 1 & 2 (ANT1/2) that have been identified^44^. While no structures of these proteins in humans have been resolved, the bovine ANT1 is available^45^ (PDB ID: 2C3E). We used this to check the validity of the predicted AlphaFold2^38^ structure, and when aligned with the bovine structure the RMSD was measured at 0.339 Å (1388 of 1906 atoms, after 5 cycles of refinement). This provided confidence that the structures would provide a good representation for these proteins. The initiator methionine was removed prior to simulation.

Another important complex identified within the inner membrane is the translocase of the inner membrane (TIM) which is found as two distinct protein complexes (TIM22 and TIM23)^46^. The TIM22 complex has been reported in humans^47^ (PDB ID: 7CGP) but had regions of protein that were not resolved. These regions were again altered using AlphaFold^38^ structures. Yeast TIM23^14^ (PDB ID: 8SCX) was used as a guide to the accuracy of a AlphaFold2 multimer^48,49^ prediction for the human structure. This is the structure that differs the most (Figure 1B) with the sequence identity for the chains in the protein complex ranging from 27-49%, leading to the observed structural differences. This highlights the possibility to improve this model as more experimental data emerges.

Working in conjunction with the TIM complex, the outer mitochondria membrane houses the translocase of the outer membrane (TOM)^46^ which is comprised of multiple copies of 5 protein chains. The structure of this has been solved from humans^15^ (PDB ID: 7VD2) but multiple chains were missing residues. To solve these issues, the protocol used for the respiratory chain complex was implemented to complete the structure. The residues added belonged mainly to disordered regions, explaining the lack of density resolved experimentally (Figure 1C). The areas of the structure that did overlap with the solved structure provided an RMSD of 0.758 Å (2947 of 3691 atoms, after 5 cycles of refinement). Again, this provided enough confidence in this model to continue.

After a protein has been translocated into the mitochondria, if it is an outer membrane β-barrel the sorting and assembly machinery (SAM) will aid in the insertion into the OMM^16,46^. The structure of this complex has been resolved in yeast^16^ (PDB ID: 7E4H) which was used as a template to construct the human homologue. Similarly to the TOM complex, there were large regions of disordered protein.

One of the most abundant proteins reported is voltage-dependent anion channel 1 (VDAC1)^22^. The structure of VDAC1 has been resolved in humans^50^ (PDB ID: 6TIQ) and this was used as the structure with E73 deprotonated^18,51^. While dimers and trimers have been reported^18,52,53^, the exact configuration and most dominant state is still unclear. For this reason, VDAC1 was included in its monomer state.

A well-reported yet poorly understood complex at the molecular level is the mitochondrial contact site and cristae organizing system (MICOS). This complex of 7 proteins has been reported to be critical for cristae topology and formation^3,54–56^, also forming part of the intermembrane bridging complex (IMB)^57,58^. This complex is critical to the accurate representation of a cristae junction, but there are no published structures of these proteins from any species. The complex can be split into two subcomplexes^59^, and we attempted to model them separately with AlphaFold2^38,49^ with very limited success. The smaller subcomplex (MIC10-26-27) had extremely low confidence and a very unclear transmembrane region. The larger complex (MIC60-19-25) exceeded the size limits of what was possible to predict with AlphaFold2. The release of AlphaFold3^60^ enabled the generation of a model of this larger subcomplex (Figure 2A). Regions that have been reported to contact the outer membrane and those anchored to the inner membrane were grouped together which provided confidence in the prediction. The entire MICOS complex could be modelled as folded, but the resulting structure did not align with the reported biochemical data and therefore was not included. This is another critical component of this system that should be updated as new structures can be generated or resolved. As the MIC60 subcomplex has been reported to contact the outer membrane and maintain distance between the membranes^57,58,61^, we chose to include this even as a partial complex.

**Figure 2:**
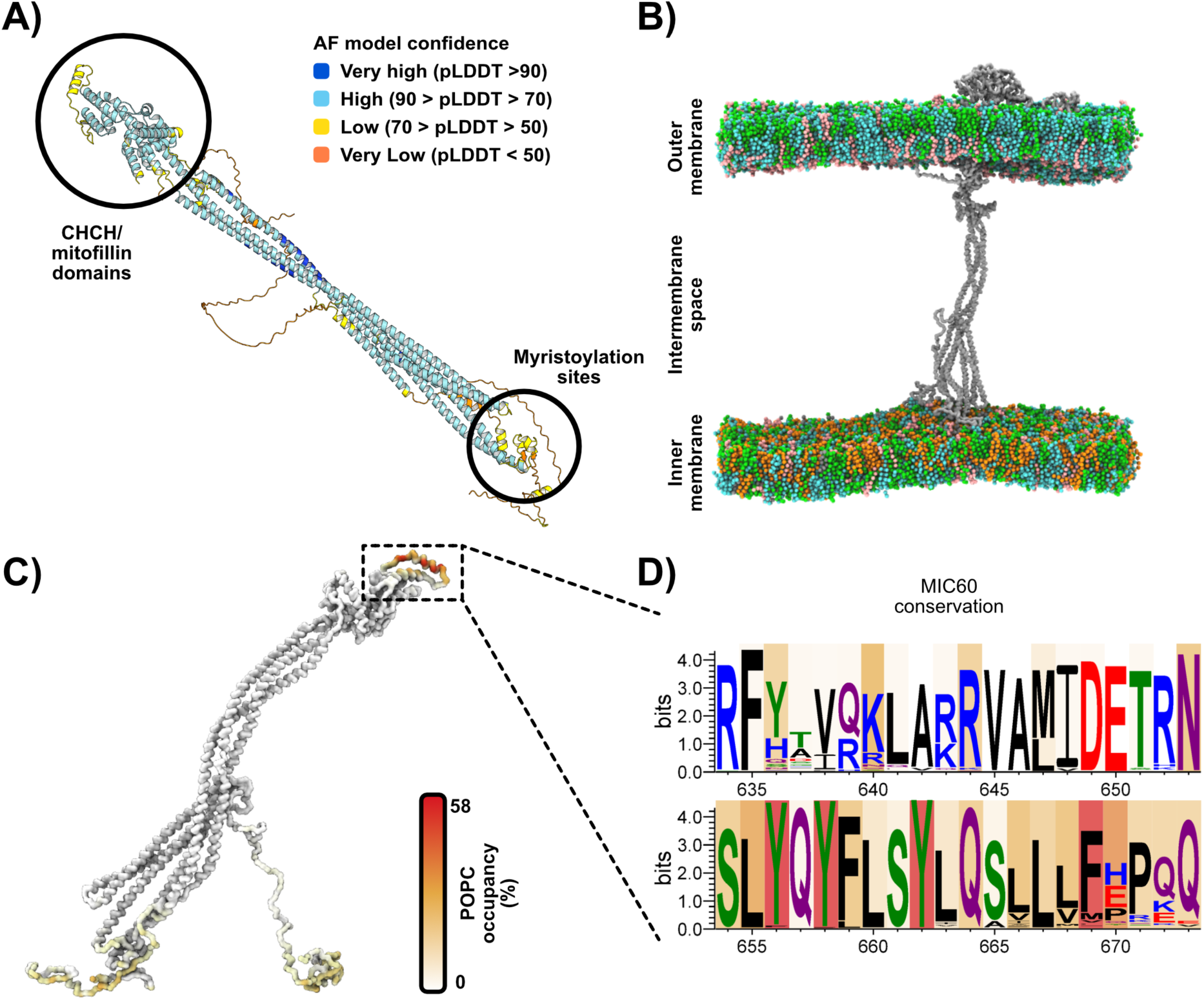
Modelling of the MIC60 subcomplex and simulation set-up. (A) The predicted structure of MIC60-19-25 complex using AlphaFold 3. The structure is coloured using the coloraf tool for PyMOL^62^, showing the confidence of the structure prediction. (B) A simulation snapshot containing the MIC60 subcomplex in the IMM and the SAM complex in the OMM. The protein is shown as a grey surface and the lipids are shown as spheres. (C) The MIC60 subcomplex colored by the contact with the headgroups POPC, the major lipid in the OMM. The color shows the amount of contact, with a darker color showing increased lipid occupancy. (D) The height of the residue shows conservation, with the color of the letters reflecting the chemistry of the residue. The background color shows the occupancy of lipids over the simulation, matching that shown in C.

### High-throughput simulations to equilibrate and analyse protein (complex) structures

After preparing the structures and assembling the protein complexes, a set of individual simulations were performed for each. This was to ensure that protein placement in the membrane was correct, any local lipid environments around the proteins could form and any membrane curvature induced by the proteins could be observed and quantified. The *martinize2*^63^ tool was used to convert the protein structures from an atomistic (AT) representation to CG, including the post-translational modifications for the MIC60 subcomplex^64^.

One complication in this process was the scale of the super complex, which was too large to convert into a CG representation as one complex. This led to converting all subunits into a CG model separately and combining the coordinates and parameter files after. To ensure individual complexes would stay coordinated throughout the simulations, an elastic network was added within complexes after these steps.

Correct orientation of the protein in the membrane is critical for meaningful simulations. We used the OPM server^65,66^ as it contains a plethora of oriented mitochondrial proteins, as well as their online server to generate any that were missing. Depending on the localization, the protein or protein complex was embedded in a membrane resembling either the inner^67^ (phosphocholine (POPC): phosphatidylethanolamine (SAPE): phosphatidylinositol (PAPI): phosphatidylserine (POPS): cardiolipin (CDL2); 29:36:6:3:26, POPC:SAPE:PAPI:POPS:CDL2; 58:37:5:3:11 in upper and lower leaflets respectively) or outer^18^ (POPC:SAPE:PAPI: cholesterol (CHOL): ceramide (PCER); 42.5:32:5:15.5:5, POPC:SAPE:PAPI:CHOL; 52:14:19:15 in upper and lower leaflets respectively) mitochondrial membrane. The workflow is shown in Figure 3A.

**Figure 3:**
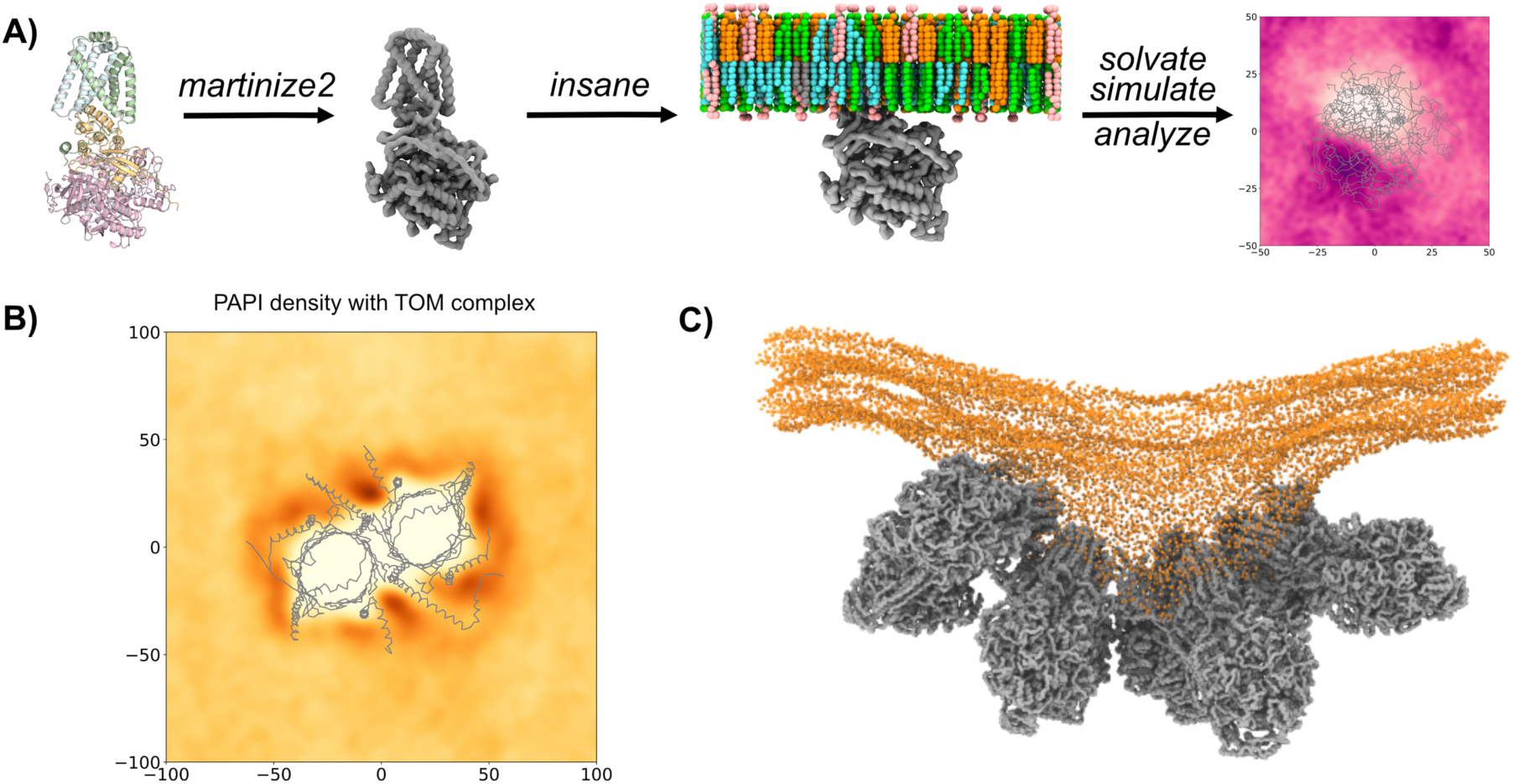
High throughput simulation set up and analysis. (A) The workflow performed in the high-throughput simulations of the selected proteins. The protein was converted to CG, embedded in the membrane and multiple independent simulations performed. These trajectories were then analyzed. (B) An example of a lipid density plot, showing PAPI density around the TOM complex. A darker color indicates higher density. (C) Membrane curvature induced by a row of ATPase dimers. The protein is shown as a grey surface, while the phosphate headgroups are shown as orange spheres.

After 5 independent repeats of short (∼1 µs) simulations, we measured the lipid density for each species present as well as the membrane deformation (Figure 3B,C and SI Figures 1-10). Hotspots of different lipid species indicate specific binding sites which can be investigated further with programmes such as PyLipID^68^ or ProLint^69^. Final coordinate positions are used in the next steps of the workflow.

Examining the lipid densities measured, we observed an enriched area of cardiolipin around ANT2 (SI Figure 5) which agrees with previous computational results^30^ but did not see the same for ANT1 (SI Figure 4). A similar accumulation of cardiolipin was seen for the transmembrane region of the ATPase dimer (SI Figure 1) again showing agreement with previous studies^70^. To fully assess these binding sites, longer simulations would need to be performed. VDAC1 has been reported to act as a phospholipid scramblase when in the dimer form^71^, and therefore the accumulation of phosphoinositol around VDAC1 (SI Figure 10) is unsurprising. In fact, most proteins showed some enrichment of PAPI when compared to bulk. Future work is needed to determine if this is an artifact of the simulations, or a biologically relevant finding.

The MIC60 subcomplex has been reported to be a member of the IMB, forming contacts with both the membrane and the C-terminal region of SAM50^57,58,61^. To test if this is observed with these models, the MIC60 subcomplex was simulated for 5 µs with the outer membrane with the SAM complex embedded. The MIC60 subcomplex was not in contact with the outer membrane at the start of the simulations, but in 2/5 simulations contact with the OMM was made, where it also made transient contacts with the SAM complex. Although this is the minority of the simulations, when contact was made it was strongly maintained (up to 8 µs when simulations were extended). A snapshot of this can be seen in Figure 2B. The area that made contact with the membrane was identified as the lipid-binding site of MIC60 experimentally^58^ (Figure 2C,D) and these residues are highly conserved. These results provide confidence that this model will replicate what could be happening *in vivo*.

To investigate the effect of rows of dimers of ATP synthases on membrane curvature, we performed a simulation with four dimers (SI Figure 11). The effect of having multiple copies was profound, and this can be seen in Figure 3C. Even though the starting configuration was a flat membrane, significant curvature was imposed showing that this dimer row would be able to maintain curvature in the model inner membrane crista.

### Building and equilibration of the outer mitochondrial membrane

To compose the outer membrane, final poses of the SAM complex, TOM complex and VDAC1 were taken from the short simulation with an annular ring of lipids (∼1 nm) as can be seen in Figure 4. The packing programme bentopy^72^ was then used to position the proteins over the membrane. This was achieved by centring the proteins at the middle of the bilayer placement and constraining the proteins to be rotated along their z axis only. The ratios of the components were based upon experimental data^21,22^. TOM20 has been measured have approximately 500 copies per µm^2^ ^21^. While SAM50 is reported to be similar to TOM20 in terms of protein complex numbers^22^, VDAC1 is 7x more abundant^22^. When these protein copy numbers are converted to a 65 x 65 nm^2^ membrane, 1 copy of the TOM20 and SAM50 complexes and 7 VDAC1 monomers would be expected. As other protein complexes are overlooked in this study, the absolute numbers were increased approximately 3-fold in this model (Table 1). The other lipid components of the system were added using insane^73^ to make the complete outer membrane (Figure 4).

**Figure 4:**
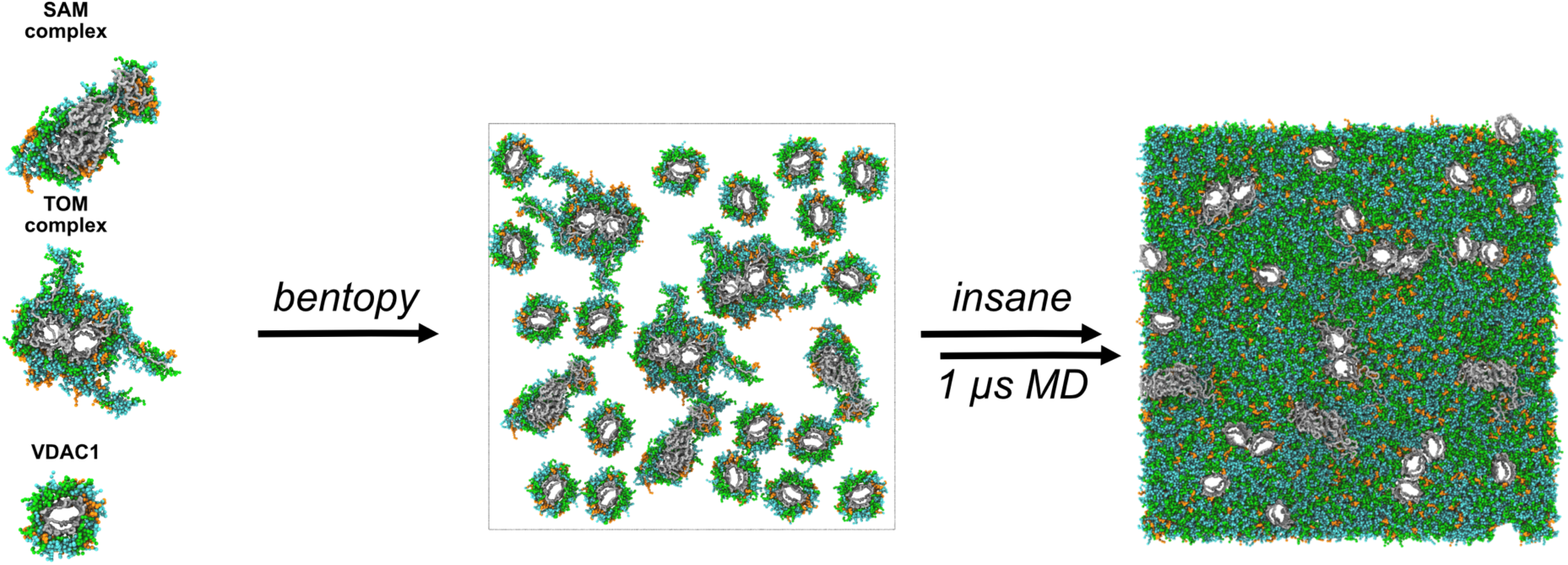
Setting up the outer mitochondrial membrane. The proteins with lipid shells (taken from the end of a simulation) were supplied to bentopy^72^, which placed a specified number of proteins with varied rotations. The membrane was then completed with lipids using the insane membrane builder and simulated for 1 µs, with the final frame shown.

The resulting model was then simulated for 1 µs. The size of the membrane stayed similar throughout the simulation, as did the volume of the simulation box and the potential energy of the system (SI Figure 11). The membrane broadly retained the flat nature, with expected undulations throughout. As the first simulation showed no great fluctuations in the system after 500 ns, the following independent repeats were simulated for this shorter time.

### Building and equilibration of the inner mitochondrial membrane

Due to the topologically challenging shape of the cristae, the classical flat bilayer construction tools could not be utilized for the IMM. In recent years, TS2CG, a method to convert a triangulated surface to a CG representation, has been developed by Pezeshkian *et al.*^31^. In this body of work, a CG representation of an entire mitochondrion (lipids only) was built. While the input density maps are available^74^, finding inividual crista that would satisfy periodic boundary conditions proved challenging. In lieu of an experimentally resolved membrane map, an artifical triangulated surface was generated using the 3D modelling software Blender. Experimental values were used as a guide to the size of the system^74,75^, resulting in a 65 nm x 65 nm bilayer in the x and y dimensions, featuring a crista of ∼20 nm wide which extended for ∼50 nm (Figure 5).

**Figure 5:**
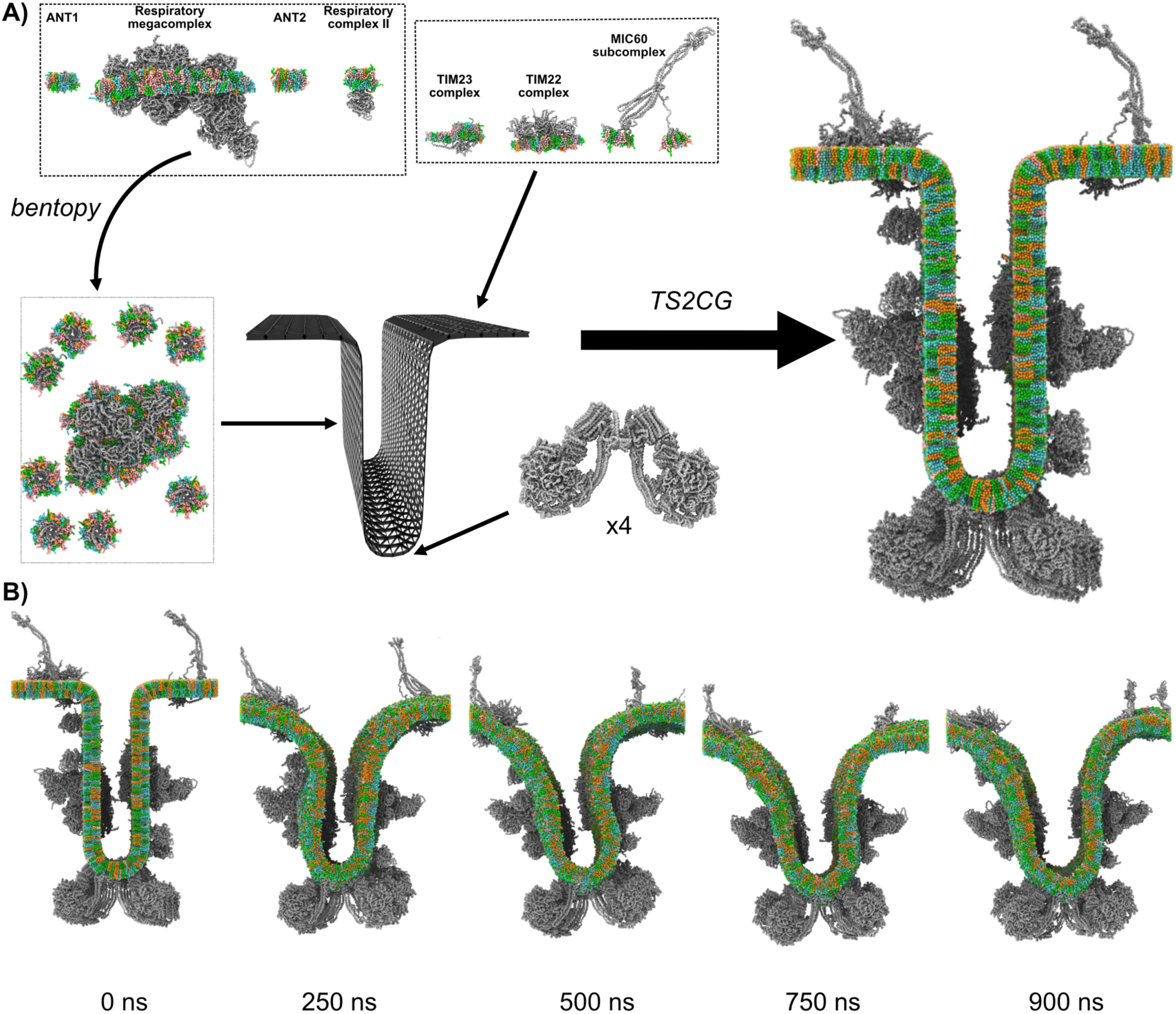
Setting up and simulating the inner mitochondrial membrane crista. A) The triangulated surface (shown as a black mesh) was constructed using Blender, resulting in a 65 nm x 65 nm bilayer in the x and y dimensions, featuring a crista of ∼20 nm wide which extended for ∼50 nm. All proteins (apart from the ATPase) were supplied with a lipid shell. They were separated according to the regions they are located in, with the crista sides being constructed using bentopy^72^. Final protein and lipid placements were performed by TS2CG, and the resultant system is shown. B) Snapshots from the simulation of the inner mitochondrial membrane.

With the triangulated surface generated, TS2CG^31^ could be used to place the proteins and the lipids found in the mitochondrial inner membrane. Single vertices were individually selected to place the proteins that are located on the inner membrane regions that face the outer membrane. This included the TIM22 complex^76,77^, the TIM23 complex^76,78^ and the MIC60 subcomplex^78,79^. To take advantage of the artificially flat cristae sides, bentopy^72^ was used to evenly distribute proteins found in these areas. The proteins placed here were the ‘Type A’ respiratory supercomplex^11,77,78,80^, respiratory complex II^80,81^ and ANT1/2^42^. The objective was to match protein ratios from literature^42^ as closely as possible. In the area per crista side (∼50 x 50 nm^2^) in this model, the copy numbers to align with prior models there would be 1 respiratory supercomplex, 1 complex II and 15 ANT monomers^42^. While the respiratory complex numbers were matched, the packing method was unable to place all copies of ANT1/2. Therefore, the stochiometry of ANT was reduced to 5-6 on each crista side.

Four ATPase dimers were placed along the ridge of the crista. Because of the high amount of curvature induced by the ATPase dimers (Figure 3C) in the smaller simulations, their lipid shells were not included in the construction of the crista junction. The angle of the dimer interface matched the modelled crista ridge well, giving more confidence to both the protein model and the membrane shape. Every other protein complex considered included their surrounding lipid shells generated from the high-throughput simulations performed previously.

Following solvation and energy minimisation, this membrane was simulated for 1 µs to check overall system stability and any changes in morphology. During the simulation the extreme curves at the membrane invagination relaxed (Figure 6A), but the angle at the tip of the crista stays similar. This suggests that the ATPase dimers maintain the modelled curvature of the crista edges, while the other curves in the system are not as tightly controlled by proteins included in this model.

**Figure 6:**
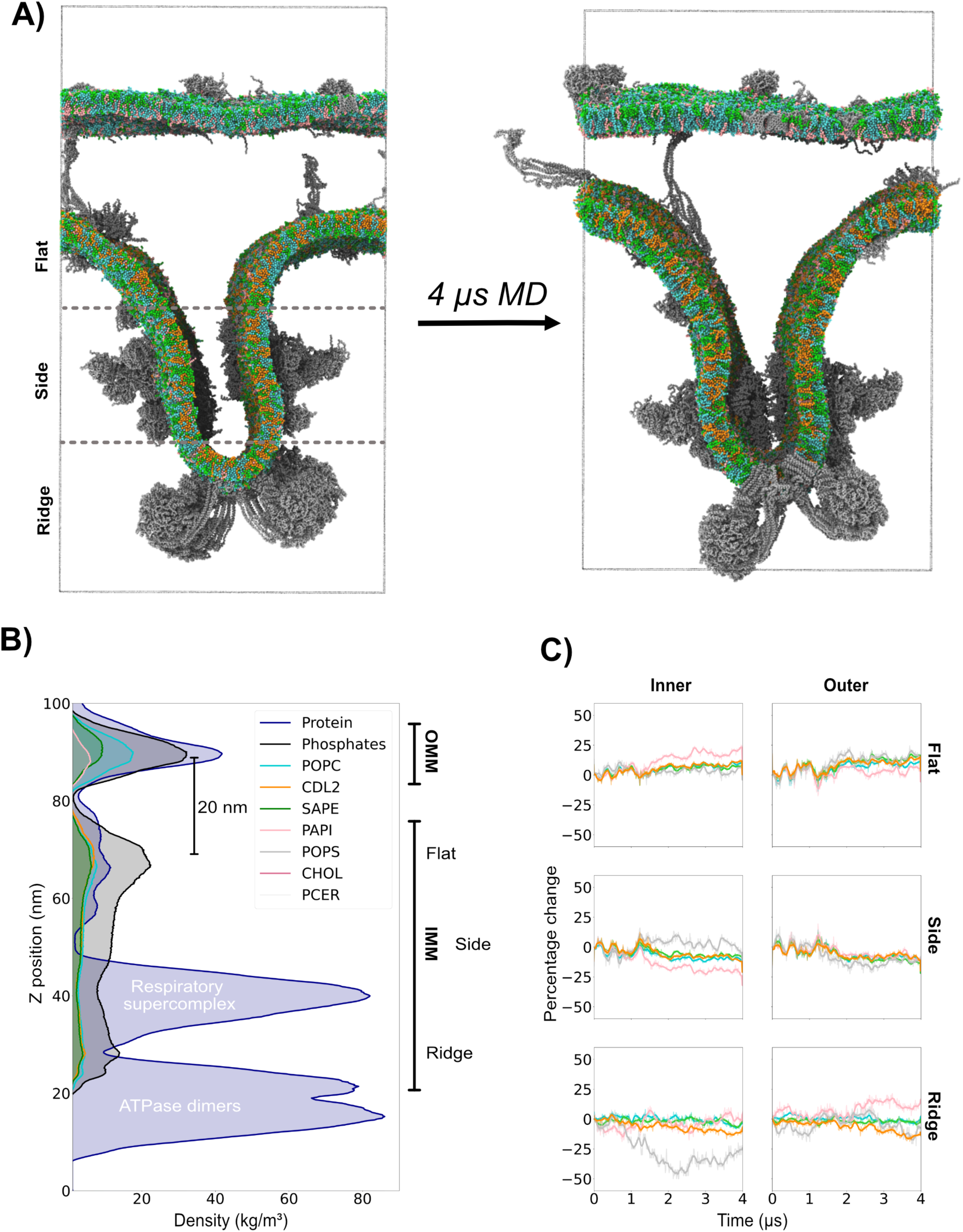
The combined membrane mitochondrial crista setup & analysis. (A) The starting and final configuration of the crista junction model. The proteins are shown in gray, while the lipids are colored according to the panel in C. (B) The density of phosphates (black) and protein backbone (dark blue) over the z-dimension of the simulation box. The positions of phosphate headgroups of POPC (cyan), CDL2 (orange), SAPE (green), PAPI (pink), POPS (light gray), CHOL (dark red) and PCER (off white) are also shown. Different regions of the system are highlighted. (C) The percentage change of lipid amounts of the IMM leaflets (shown in A) over the course of the simulation, averaged every 10 steps. The colors are the same as indicated in B.

The distance between the two edges of the crista increases over the course of the simulation (Figure 5B). This could be due to the artificial set up, but also reflective of the fact that important proteins are still missing from the system. The opening width at the cristae junctions has been proposed to be regulated in part by the long isoform of OPA1^5^. While the short isoform has been reported and studied^82,83^, the structure of l-OPA1 remains unresolved. The lack of this within the model could contribute to this widening of the opening.

### Combining the membranes into a full model of a mitochondrial crista

To combine the membranes, only the lipid and the protein components from the set up simulations were considered (i.e. solvent was removed). When assembling such a system, special attention needs to be given to the size of the membranes after the lipids have ‘relaxed’ in their previous simulations. If this is not considered, artificial membrane buckling could be induced.

The membranes were placed ∼20 nm apart to capture the width of the inner membrane space from experimental measurements^75^, ensuring that there were no overlapping atoms, especially between the MIC60 subcomplex and the lipids/proteins of the OMM. The distance between the membranes in the z-direction needed to be sufficient to ensure that any membrane deformations would not lead to membrane contact between the IMM and OMM. The overall dimensions of the resulting system were 60 x 60 x 115 nm^3^. When assembled and re-solvated, the system contains nearly 4 million particles with the composition found in Table 1.

Simulation of this system for 4 µs showed contact between the two respiratory complexes (Figure 6A) which resulted in narrowing of the crista width. It is difficult to know if this is an artifact of the simulation setup or if this could play a role in maintaining cristae membrane separation *in vivo*. The distance between the membranes were ∼20 nm (Figure 6B) with undulations which can be seen in Figure 6A. The majority of the protein density seen in the system reflects the respiratory super complex and the ATPase dimers. The dimensions of the bilayers do not change substantially over the simulation (SI Figure 13).

To get a better understanding of the movement and interactions of the proteins and lipids present in the system, several means of analysis can be used. One example of this, the change in lipid composition in regions of the inner membrane over the course of the simulation, can be seen in Figure 6C. POPS was depleted in the inner membrane at the ridge region. As the ridge of the crista is negatively curved, the reduction of this lipid aligns with previous results indicating phosphatidylserine’s positive intrinsic curvature^84,85^. As the relative number of POPS lipids is significantly lower than others in the membrane (Table 1), it is not clear which lipids replace those that are displaced. The flat region as seen in Figure 6A, includes the positive curvature of the crista junction, and in the inner membrane PAPI was enriched. This result is unusual as PAPI is reported to have negative intrinsic curvature^85,86^. As CDL2 is so important in maintaining the highly curved nature of the IMM, it is surprising to see that there is not enrichment at the crista ridge to align with cardiolipin’s intrinsic negative curvature^85^. The proteins present in the system will complicate the landscape of lipid sorting, which could be the reason that the results differ from those measured in lipid only systems^8,85^.

### Building a more realistic system separating matrix from cytoplasmic space

The mitochondrial model presented here (shown in Figure 6A) is the most accurate model produced to date. However, it is not a sufficient model for a real mitochondrion as if periodic boundary conditions are considered there is no real separation between the space that represents the matrix (inside of the IMM) and the cytosolic space (outside of the OMM). This means, with the current model, if soluble mitochondrial components are included, they will not be confined in the biologically relevant regions.

One way to overcome this issue would be to include another IMM and OMM within the model, as shown in Figure 7. This ‘MitoSlice model’ would allow solvent and metabolites diffuse through the β-barrels present in the OMM, while preventing any passage through the IMM. This will allow the inclusion of mtDNA, soluble proteins, metabolites and differing ion concentrations between the subcellular compartments. While this could be achieved with specific boundary conditions, a dual crista system is more biologically relevant.

**Figure 7:**
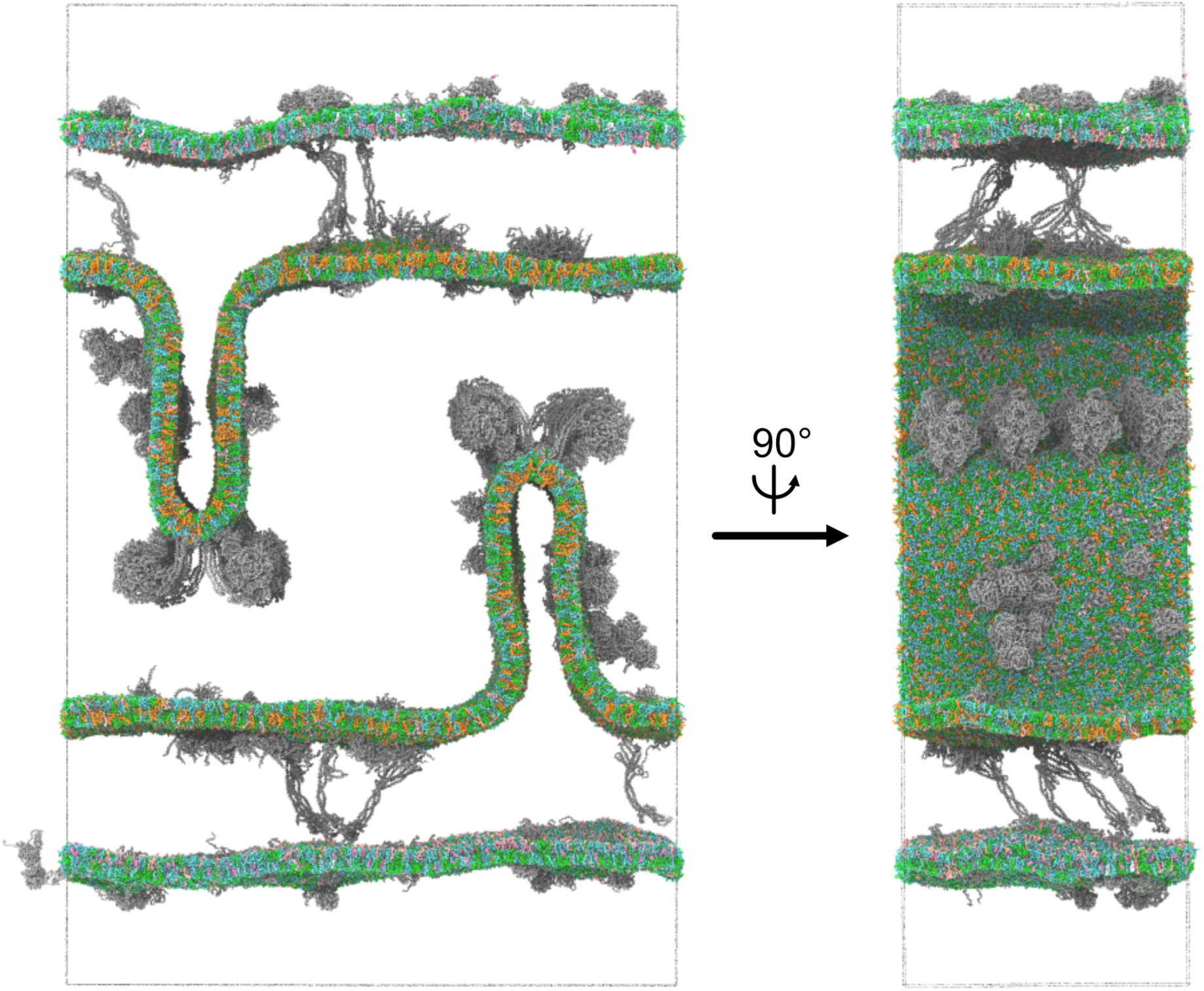
A more representative model of the mitochondrial cristae. Proteins are shown as grey surfaces.

As well as increasing the complexity through separate compartments and composition, to make the model closer to an equilibrium state, lipids could be placed based on their curvature preference^8,85,87^. Furthermore, the current model lacks the inclusion of various cofactors such as ATP, ubiquinone/ubiquinol and heme which are required for the respiratory complex function^11^. There is an ongoing development of cofactor topologies compatible with the latest version of the Martini force field, enabling their inclusion in future versions of our mitochondrial models^88,89^.

Building up the complexity does increase the size of the system, which can be seen in Table 1. The required computational resources needed to simulate systems of this size (and larger) are considerable, but it will intrinsically provide improved statistics for properties such as protein-protein interactions. Nonetheless, this draws even more importance to starting the system as close to equilibrium state as possible, to reduce the needed simulation time. As systems get larger and more complex, it also increases the difficulty of preforming analysis. Many of the existing analysis methods, especially for lipids, rely on flat or spherical bilayers. A balance between the complexity and accuracy of the simulation and the meaningful data that can be obtained should be managed.

## Conclusions

In order to study the dynamic interactions present in mitochondrial cristae, we developed the most representative system to date using integrative modelling. Particular focus was on including not only the most relevant proteins in the correct oligomerization state but ensuring all were from the same species and included all residues that would be found *in vivo*. This led to a range of approaches to including accurate protein structures, from using structures found in the PDB to verifying novel protein complex structures predicted by AlphaFold. When predicted structures could be compared to experimentally resolved structures from homologues species, the alignment was overall extremely high (on average < 1 Å), showing the quality that this approach can have.

High-throughput simulations of each of these proteins ensured that the structures would adopt an optimum position embedded in the membrane. It also provided the chance for investigation of the lipid density around the proteins and any curvature induced in the bilayer. If further specific questions about these proteins arise in the future, these simulations can be used as a resource and re-analysed or extended.

Performing the smaller scale simulations ensured the proteins could be included with an annular lipid shell, ensuring that the systems will start closer to an equilibrium state. While human mitochondria are well characterized in terms of their absolute protein copy numbers^22^ and relative stoichiometries^42^, these values proved difficult to reach even with an optimised packing algorithm^72^ when assembling each membrane. This issue is amplified by the inclusion of the local lipid shell and disordered regions of the protein, which occupy a large amount of space. It could also reflect protein densities differing at various regions throughout the cristae, which is difficult to reflect in a system such as this. As more is understood about the local arrangements of these proteins, the systems can be improved.

The inner membrane shape in this model was generated artificially, matching measurements made experimentally^75^. While this made it possible to easily place proteins and satisfy periodic boundary conditions, ideally experimental maps of the membranes will be used to harness the vast amount of data available on this matter. In the simulations containing just the IMM, opening of the crista junction was observed in all independent repeats. This indicates the importance of including all necessary proteins and the role of the OMM.

When the crista junction model was simulated, the curvature in the system was well maintained and the membranes stayed at a constant distance apart, even without the use of constraints. As the system is modest in size given the complexity, timescales on which protein diffusion and interaction can be seen were reached. However, analysis on properties such as these still poses a challenge due to the complexity and shape of the membranes included. New techniques and software will need to be developed to ensure systems such as these can be meaningfully analysed.

Finally, a more complicated system, the MitoSlice model, was presented as a method to be able to compartmentalize the soluble regions of the mitochondria. This will be crucial if components such as cytochrome c or mtDNA are to be included.

Overall, this is the most comprehensive model of a crista produced to date, including the most relevant proteins/complexes and both membranes with their realistic lipid composition. However, there are still many membrane proteins missing and a complete lack of soluble components. This work provides a workflow for increasing the complexity of this system and is intended to be a ‘living model’. Building up this model gradually provides opportunity not only to ensure that each component is correct, but to allow development of methods and tools to build and analyse these systems. Collective input of knowledge and methodology from the community on these issues will help us to reach the goal of creating a model of an entire mitochondrion.

## Methods

### Preparation of protein structures

As the aim was to achieve an accurate model, all residues that are present in the transcribed sequence excluding signal peptides were included. Refinement of each structure was a manual process which relied on user intuition and knowledge. When preparing the structures for each of the proteins or complexes, four different methods were utilized: Using the PDB file (VDAC1), using the PDB file but altering the oligomeric state (ATPase), using a template PDB file and aligning AlphaFold monomers (Respiratory super complex, complex II, TIM22, TOM) or using AlphaFold to model the overall structure (ANT1/2, TIM23, SAM, MIC60). When preparing all structures, any non-protein entities were removed from the structure.

#### Using the PDB file

The PDB file for VDAC1 was downloaded from the database (PDB ID: 6TIQ) and inspected to ensure there were no missing residues. As previous work has shown the importance of the residue E73 being deprotonated^71,90^ it was included.

#### Using the PDB file but altering the oligomeric state

The PDB file of the human ATPase monomer (PDB ID: 8H9S) was duplicated and aligned with each ATPase monomer from the bovine structure (PDB ID: 7AJB) using PyMOL.

#### Using a template PDB file and aligning AlphaFold monomers

This method relied upon finding a template structure file for each complex (Respiratory super complex PDB ID: 8UGH, Complex II PDB ID: 8GS8, TIM22 PDB ID: 7CGP and TOM PDB ID: 7VD2).

When the template structure was not from a human, the homologous chain from humans was identified using UniProt^43^. Using UniProt IDs, the relevant chains were then downloaded from the AlphaFold2 database^38^. Using the PyMOL align command, each chain was then aligned with the relevant subunit from the template structure. Any subunits that did not align well were not included in the model. As AlphaFold models all residues transcribed for a protein, UniProt^43^ was used to check if there were any signal sequences present, as these would be removed *in vivo*. When they were identified, the residues were removed using PyMOL. Then, all subunits were saved as one complex and critically assessed to ensure there were no overlapping atoms. To assess the final quality of each of these newly produced structures, the new structure was aligned with the template in PyMOL and RMSD values (after 5 cycles of refinement) were reported.

#### Using AlphaFold to model the overall structure

When appropriate templates were not available, AlphaFold^38,48,49,60^ was utilized to generate the complete structures. This followed the stoichiometries and oligomerization states found in literature for each of these complexes. AlphaFold2^38,48^ and ColabFold^49^ was used for all proteins previously listed apart from MIC60 which used AlphaFold 3^60^. Following prediction, as previously described, signal peptides were removed.

The sequence identity for the TIM23 complex (TIM17, TIM23 and TIM44) was done by comparing the sequences from Homo sapiens and Saccharomyces cerevisiae using the NCBI BLAST server.

### High-throughput simulations to equilibrate and analyse protein (complex) structures

To correctly orientate the proteins within the mitochondrial membrane, the PPM web server^65,66^ was used to centre the transmembrane region of the protein for use in the following steps. The dummy beads provided as membrane positions were removed from the structure files. To convert the proteins into a CG representation, martinize2^63^ was used to generate the coordinate and itp files. The Martini 3^36^ force field was used, with an elastic network (500 kJ mol^-1^ nm^-2^) between all chains in a given structure.

The only exception to this is the respiratory super complex, which proved too large to use the martinize2 as one complex. To overcome this, the subunits were converted separately and then combined into one structure and itp file per complex (i.e. Complex I, complex III and complex IV). The inter-subunit elastic network was then constructed by identifying backbone particles that were within 1 nm of each other, and construct bonds for these. Again, these were given a force constant of 500 kJ/mol.

The correctly positioned CG proteins were then fed into the insane tool^73^ to construct the membrane. Depending on the protein, they were place in either inner^67^ (POPC:SAPE:PAPI:POPS:CDL2; 29:36:6:3:26, POPC:SAPE:PAPI:POPS:CDL2; 58:37:5:3:11 in upper and lower leaflets respectively) or outer^18^ (POPC:SAPE:PAPI:CHOL:PCER; 42.5:32:5:15.5:5, POPC:SAPE:PAPI:CHOL; 52:14:19:15 in upper and lower leaflets respectively) lipid compositions. A hexagonal simulation box was used to reduce simulation time needed, and the system was solvated with Martini water. Ions were added to the system with *gmx genion*, with 150 mM of NaCl being used to neutralise the system. The simulation box was then energy minimized using the steepest descents algorithm for 5000 steps.

Production simulations were performed in an NpT ensemble at 310 K and 1 bar for 1 µs, using a timestep of 20 fs. Temperature coupling was performed with the Velocity rescaling algorithm^91^, with protein, lipids and solvent & ions coupled separately. The temperature coupling constant was set at 1 ps. The Parrinello-Rahman algorithm^92^ was used for pressure coupling with a τ*_p_*= 12.0 ps in a semi-isotropic manner. A single cut-off of 1.13 nm was used for Van der Waals interactions and the reaction-field algorithm^93^ was used for electrostatic interactions with a cut-off 1.1 nm^94^. Coordinates were saved every 1 ns. 5 independent simulations were performed for each protein. All simulations were performed with GROMACS 2021.3^95,96^.

When investigating the effect of multiple ATPase dimers on membrane curvature, a row was artificially created by arranging four dimers in a row in PyMOL. An elastic network was used to hold monomers in dimers in the correct orientation, but not used between dimers. This was then embedded in a bilayer and simulated for 5 µs.

Simulations with MIC60 in the IMM and SAM in the OMM were constructed by assembling the membranes separately and then modifying the z-position of the OMM using *gmx editconf*. The membranes were then saved into one coordinate file and solvated. The rest of the simulation workflow was the same as described. All 5 simulations were run for 5 µs, with one being extended to 8 µs. PyLipID^68^ was used to measure the contacts between MIC60 and POPC. Conservation of the residues was produced by the Consurf server^97^ and the logo generated with the WebLogo3 server^98^.

Analysis of lipid density was performed using an in-house analysis script. Briefly, all trajectories were aligned on the protein and density of the phosphate heads plotted using matplotlib^99^. Other visualization of the systems were performed using VMD^100^.

### Building and equilibration of the outer mitochondrial membrane

VDAC1, SAM and TOM structures were taken from a final pose of the small simulation, with an annular lipid shell. This was obtained using PyMOL and selecting lipids within 10 nm of the protein. These structure files were then provided to bentopy^72^ to arrange within a 65 x 65 nm space. The rotation for each protein was restricted to just the z-direction, and rules for placing reflected the lipid band being in the z-center of the simulation box (which varied for each protein). The copy number for each protein reflected values from literature as is discussed in the main text and shown in Table 1.

When the proteins were placed, lipids were added in using the insane tool^73^ with the - ring flag to ensure lipids between proteins. This also placed lipids inside of β-barrel proteins and these were manually deleted using PyMOL. The system was then solvated, and ions added as previously described. After energy minimization, one independent simulation was performed for 1 µs, with two others running for 500 ns. The other simulation settings are identical to that for the high-throughput simulations. Analysis performed on this system was done using the *gmx energy* command, and plotted using matplotlib^99^.

### Building and equilibration of the inner mitochondrial membrane

The 3D modelling suite Blender was used to create the membrane shape based on experimentally determined values for size. This was saved as an .obj file, then converted to a .tsi file using an in-house script. The proteins to place on the crista sides were placed using bentopy^72^ as previously described but with different protein structure inputs. These were placed on the centre vertex of the crista side. The row of ATPase dimers was placed using the middle vertex of the crista ridge. For the proteins placed on the flat upper region of the system, an assortment of vertices were selected. TS2CG^31^ was used to convert this into a system containing the placed proteins and lipids. Lipids that were within 0.3 nm of each other were removed. The system was solvated, ions added and energy minimised as described. The system was simulated for just under 1 µs for the three independent repeats, with other simulation settings the same as discussed. The system was analysed in the same way as the outer membrane simulations.

### Combining the membranes into a full model of a mitochondrial crista

The membranes were combined, paying particular attention to the size of the membranes to ensure there was not a mismatch. This was achieved by using the *gmx editconf* command to move the OMM 20 nm away from the IMM, making sure that the MIC60 particles were not overlapping with the outer membrane. The system was then solvated, ions added and energy minimised as previously discussed. Production simulations were performed with the same settings detailed for the high-throughput systems. The system was simulated for 4 µs.

To identify the separate leaflets of the system, MDVoxelSegmentation^101^ was used. This information was then combined with MDAnalysis^102,103^ to measure the number of lipids in each segment where the ridge was z < 250 nm, the side 250 nm < z > 500 nm and the flat region above this. The density of the simulation components were measured with *gmx density*, discarding the first 1 µs of simulation time. Plots were created using matplotlib^99^.

The larger ‘MitoSlice’ model was created using the same method as described for the single crista but using more membranes. This system was simulated for 200 ns to check stability.

