## Supplemental information for "An integrative modelling approach to the mitochondrial cristae"

**SI Table 1**

| Protein (complex) | Original structure (PDB ID): organism | Method used in this study |
|---|---|---|
| ATPase | 8H9S: Human (overlayed with 7AJB: Bovine) | Overlaid human monomer on bovine dimer |
| Respiratory supercomplex | 8UGH: Porcine | AlphaFold 2 replacement |
| Respiratory complex II | 8GS8: Human | AlphaFold 2 replacement |
| ANT1 | 2C3E: Bovine | AlphaFold 2 Database |
| ANT2 | N/A | AlphaFold 2 Database |
| MIC60 subcomplex | N/A | AlphaFold 3 online server |
| TIM22 complex | 7CGP: Human | AlphaFold 2 replacement |
| TIM23 complex | 8SCX: Yeast | AlphaFold 2 online server |
| TOM complex | 7VD2: Human | AlphaFold 2 replacement |
| SAM complex | 7E4H: Yeast | AlphaFold 2 replacement |
| VDAC1 | 6TIQ: Human | Use original structure |

A table showing the protein (complexes) used in this study, the structure the model was based upon or compared to and then the method used to construct the model.

**SI Figure 1**

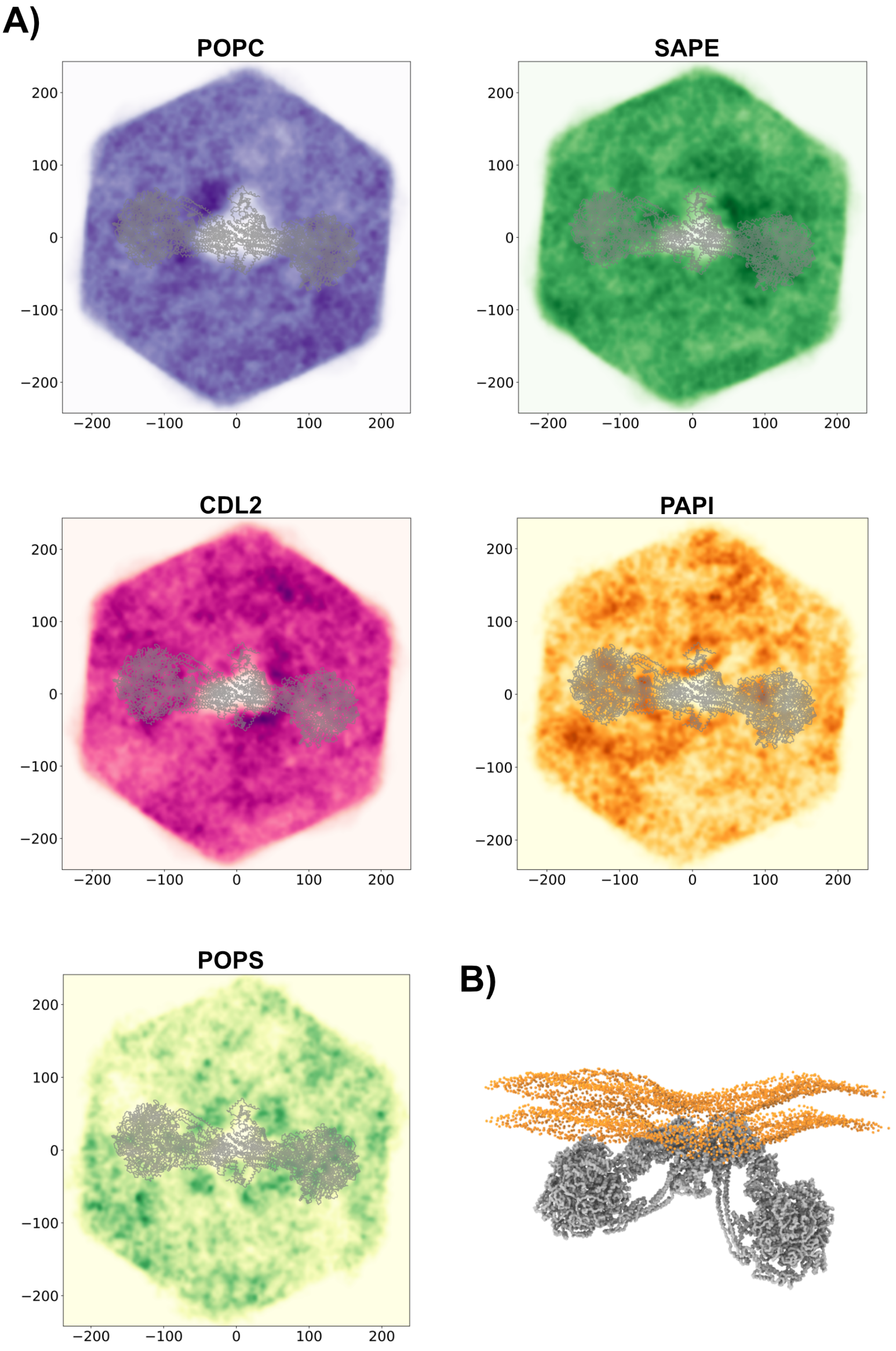


ATPase high-throughput results. A) Lipid density plots for each lipid in the system. The darker the color indicates the higher the density over all simulations. The protein backbone is shown as gray lines. B) A final frame from a simulation, showing the effect on membrane curvature. The phosphate headgroups are shown as orange spheres and the protein backbone as a gray surface.

**SI Figure 2**

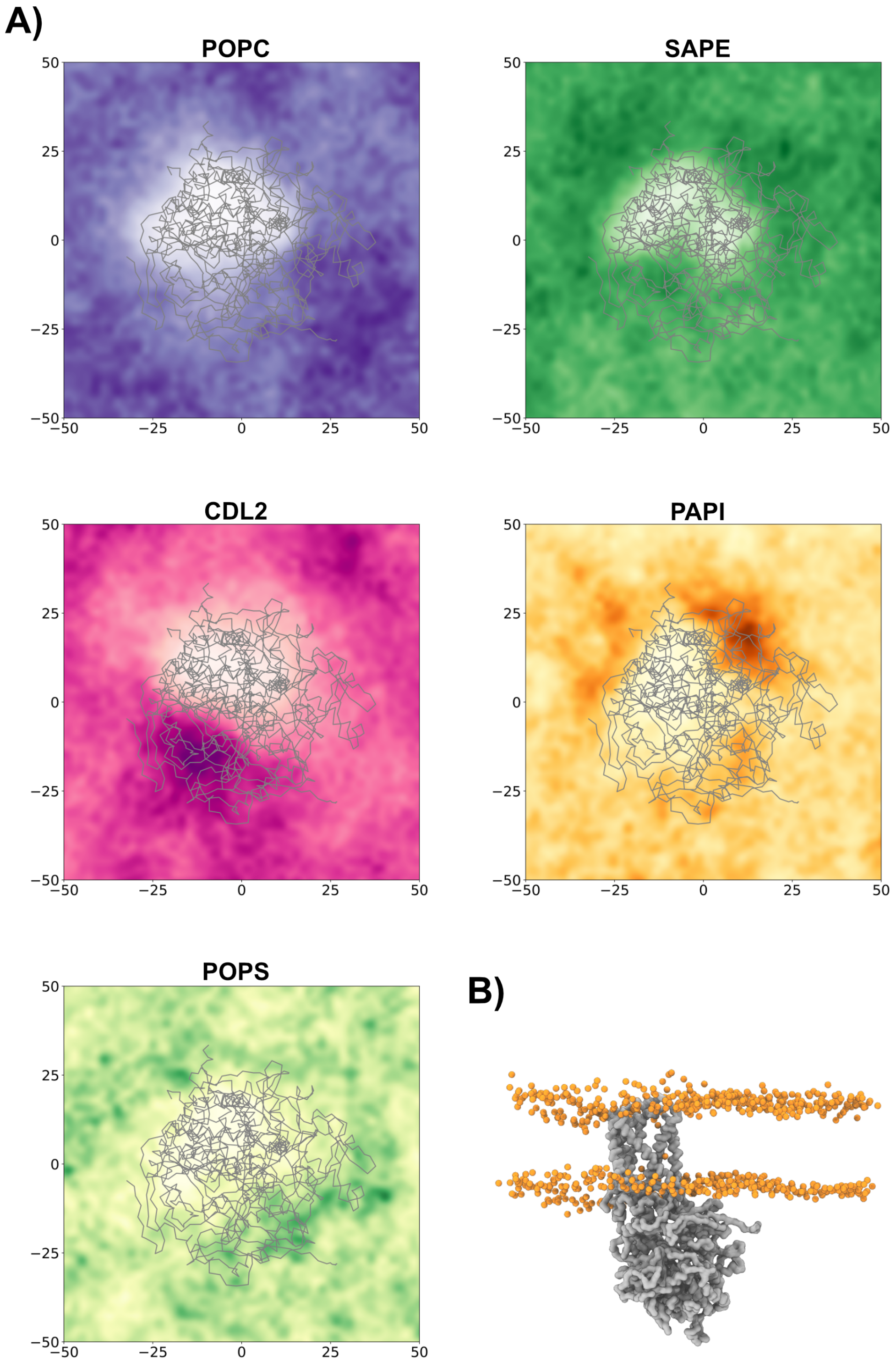


Complex II high-throughput results. A) Lipid density plots for each lipid in the system. The darker the color indicates the higher the density over all simulations. The protein backbone is shown as gray lines. B) A final frame from a simulation, showing the effect on membrane curvature. The phosphate headgroups are shown as orange spheres and the protein backbone as a gray surface.

**SI Figure 3**

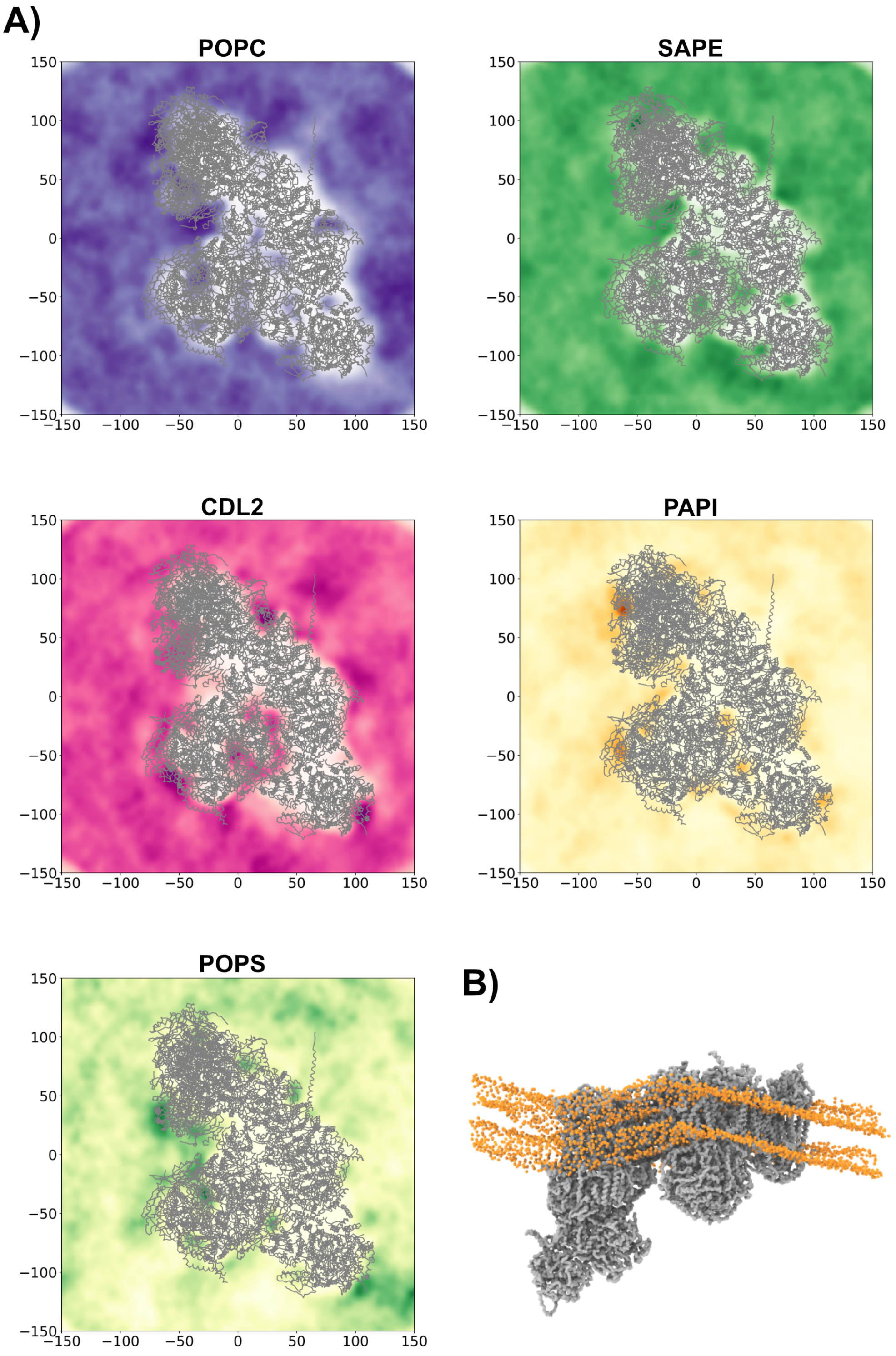


Respiratory supercomplex high-throughput results. A) Lipid density plots for each lipid in the system. The darker the color indicates the higher the density over all simulations. The protein backbone is shown as gray lines. B) A final frame from a simulation, showing the effect on membrane curvature. The phosphate headgroups are shown as orange spheres and the protein backbone as a gray surface.

**SI Figure 4**

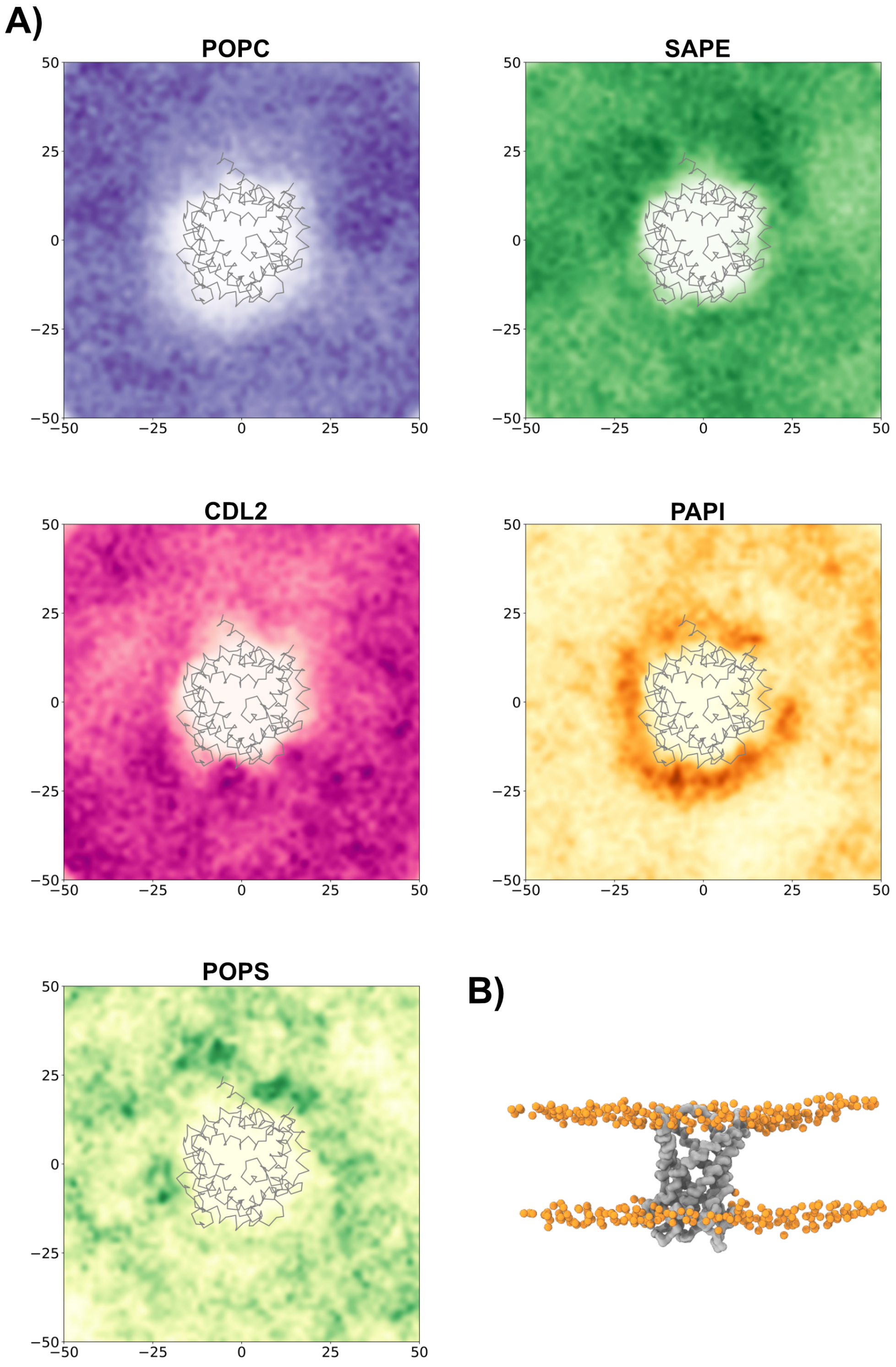


ANT1 high-throughput results. A) Lipid density plots for each lipid in the system. The darker the color indicates the higher the density over all simulations. The protein backbone is shown as gray lines. B) A final frame from a simulation, showing the effect on membrane curvature. The phosphate headgroups are shown as orange spheres and the protein backbone as a gray surface.

**SI Figure 5**

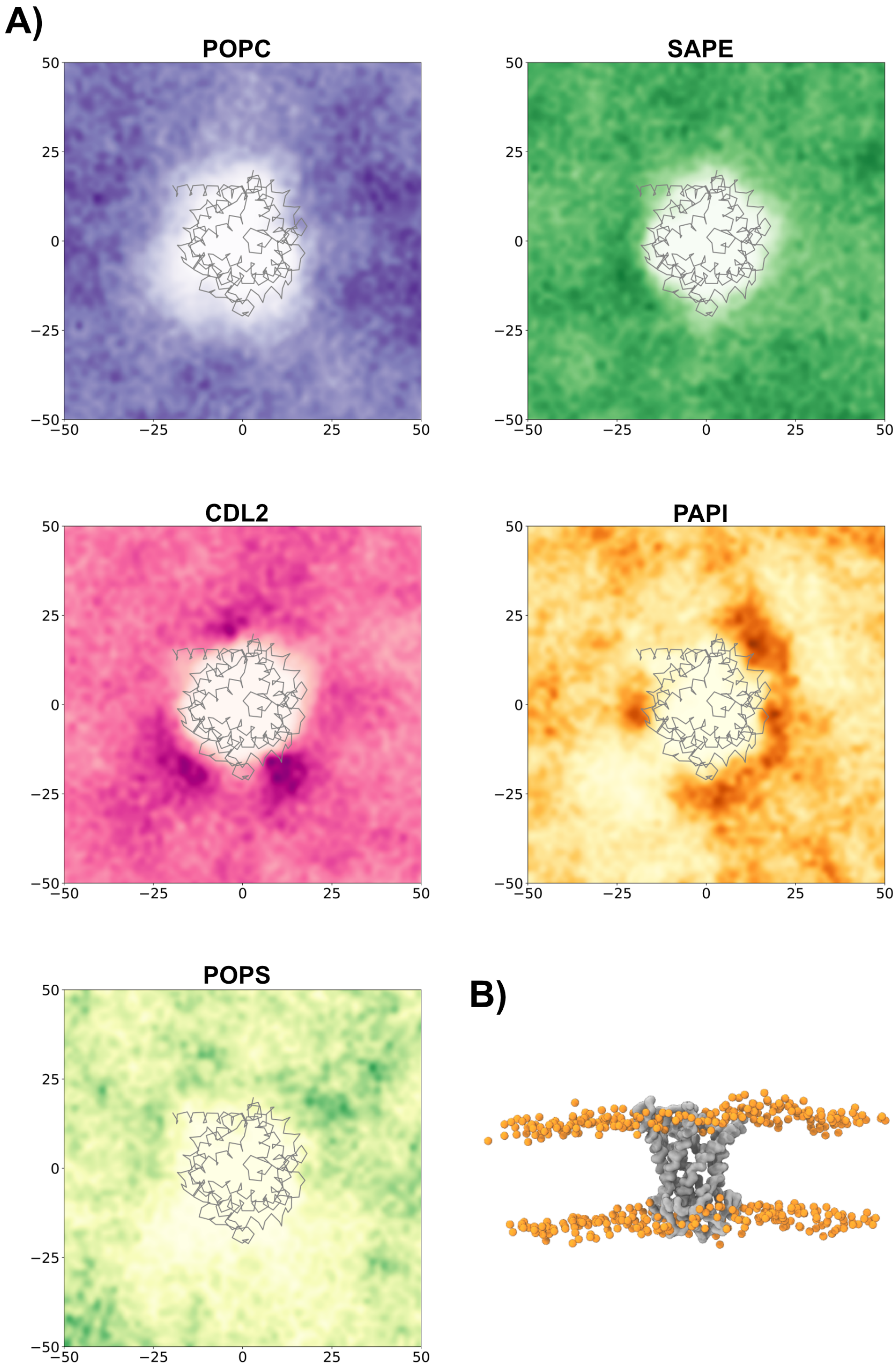


ANT2 high-throughput results. A) Lipid density plots for each lipid in the system. The darker the color indicates the higher the density over all simulations. The protein backbone is shown as gray lines. B) A final frame from a simulation, showing the effect on membrane curvature. The phosphate headgroups are shown as orange spheres and the protein backbone as a gray surface.

**SI Figure 6**

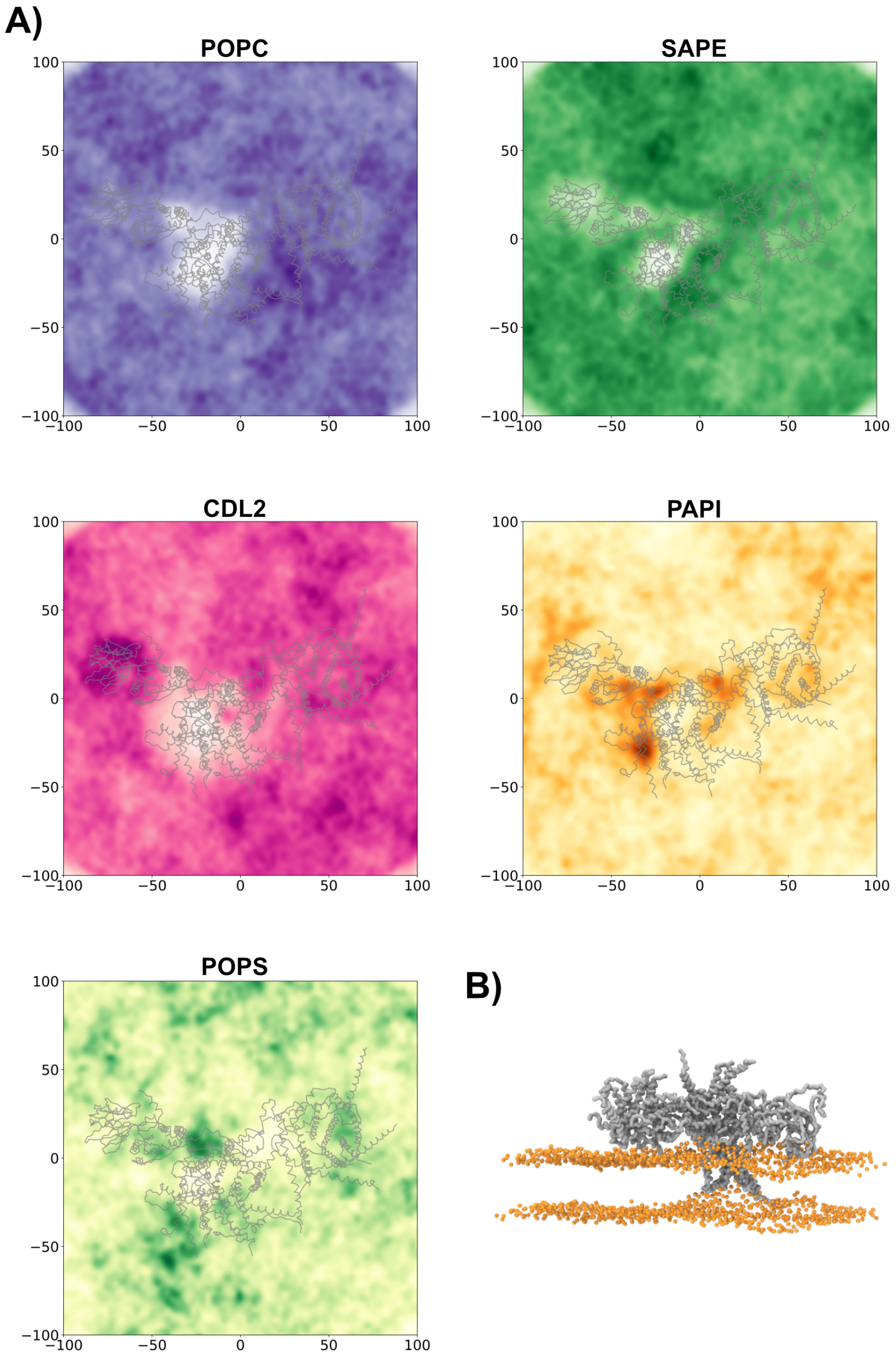


TIM22 high-throughput results. A) Lipid density plots for each lipid in the system. The darker the color indicates the higher the density over all simulations. The protein backbone is shown as gray lines. B) A final frame from a simulation, showing the effect on membrane curvature. The phosphate headgroups are shown as orange spheres and the protein backbone as a gray surface.

**SI Figure 7**

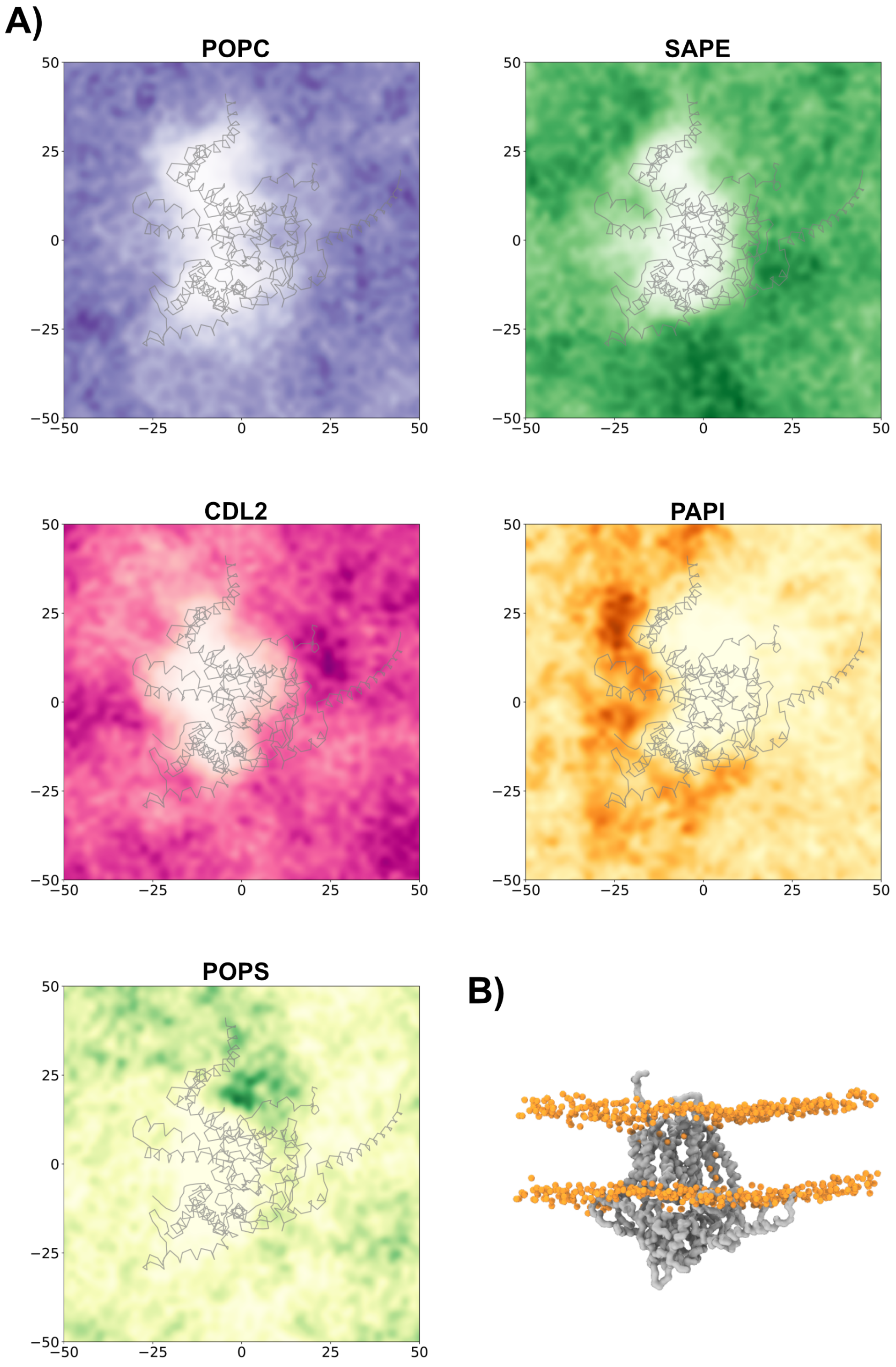


TIM23 high-throughput results. A) Lipid density plots for each lipid in the system. The darker the color indicates the higher the density over all simulations. The protein backbone is shown as gray lines. B) A final frame from a simulation, showing the effect on membrane curvature. The phosphate headgroups are shown as orange spheres and the protein backbone as a gray surface.

**SI Figure 8**

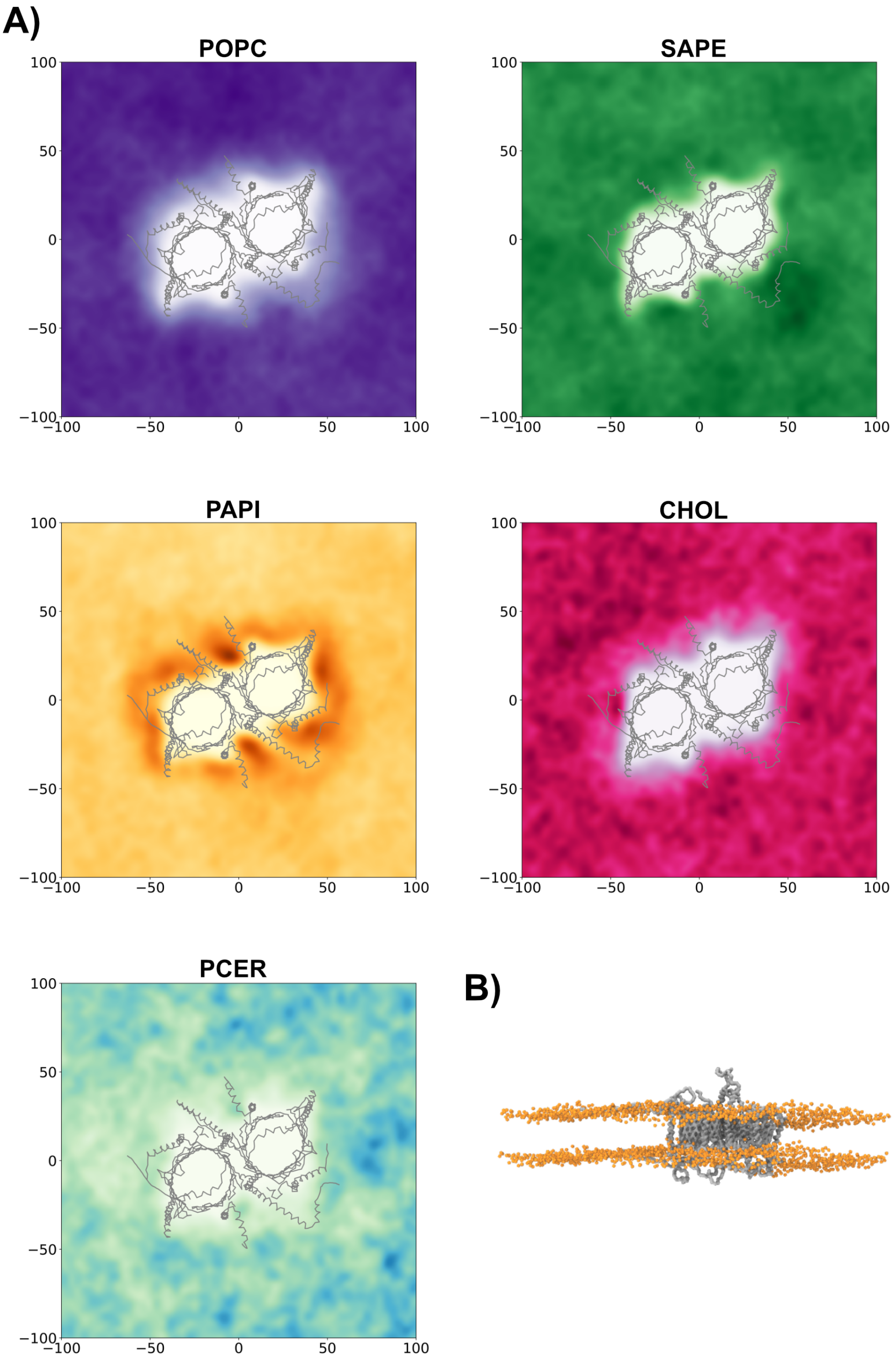


TOM complex high-throughput results. A) Lipid density plots for each lipid in the system. The darker the color indicates the higher the density over all simulations. The protein backbone is shown as gray lines. B) A final frame from a simulation, showing the effect on membrane curvature. The phosphate headgroups are shown as orange spheres and the protein backbone as a gray surface.

**SI Figure 9**

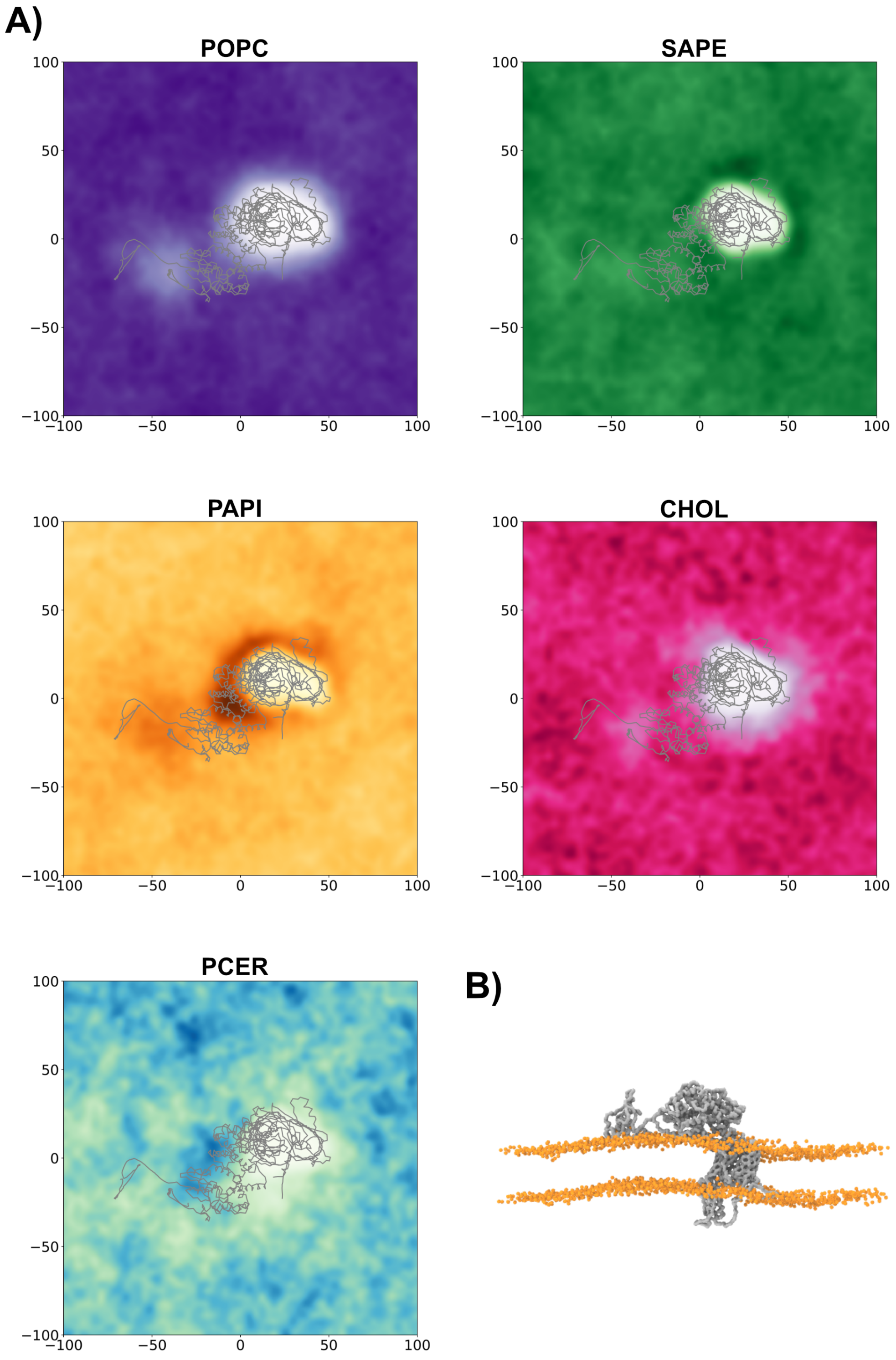


SAM complex high-throughput results. A) Lipid density plots for each lipid in the system. The darker the color indicates the higher the density over all simulations. The protein backbone is shown as gray lines. B) A final frame from a simulation, showing the effect on membrane curvature. The phosphate headgroups are shown as orange spheres and the protein backbone as a gray surface.

**SI Figure 10**

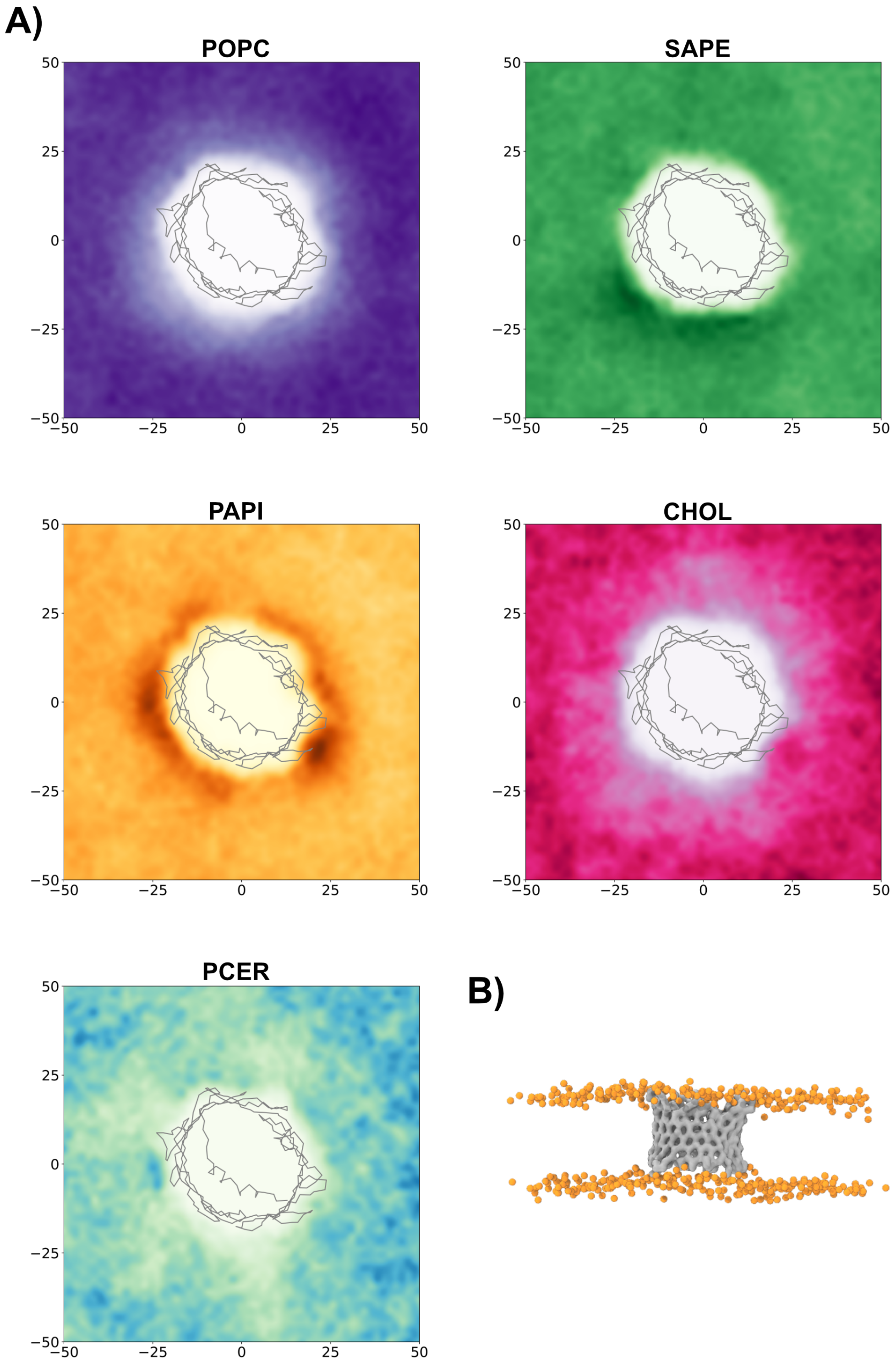


VDAC1 high-throughput results. A) Lipid density plots for each lipid in the system. The darker the color indicates the higher the density over all simulations. The protein backbone is shown as gray lines. B) A final frame from a simulation, showing the effect on membrane curvature. The phosphate headgroups are shown as orange spheres and the protein backbone as a gray surface.

**SI Figure 11**

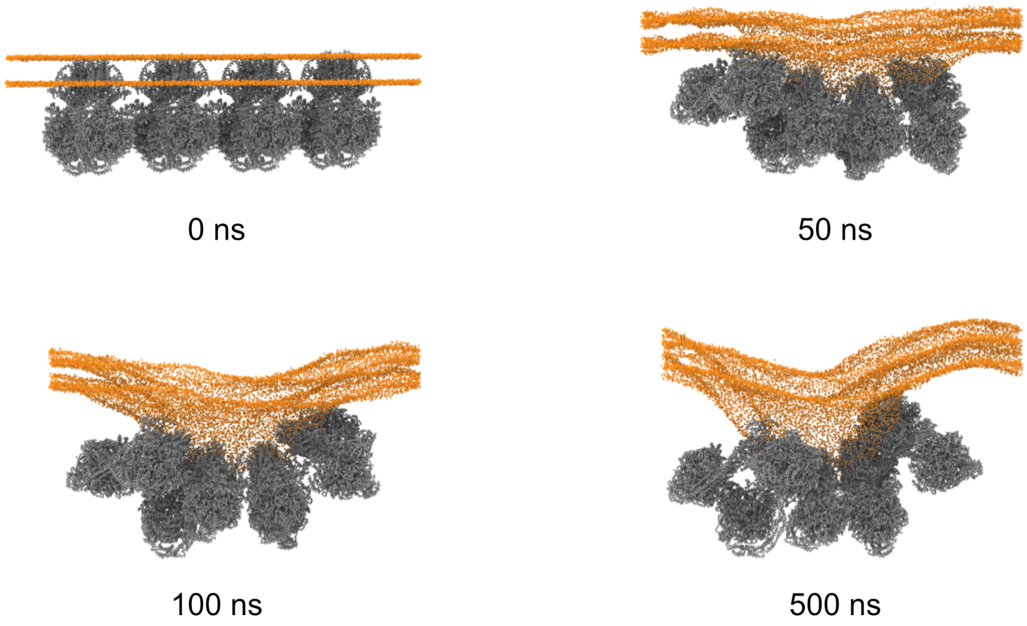


Snapshots from a simulation with a row of ATPase dimers. The phosphate head groups are shown as orange spheres and the protein backbone is shown as a gray surface.

**SI Figure 12**

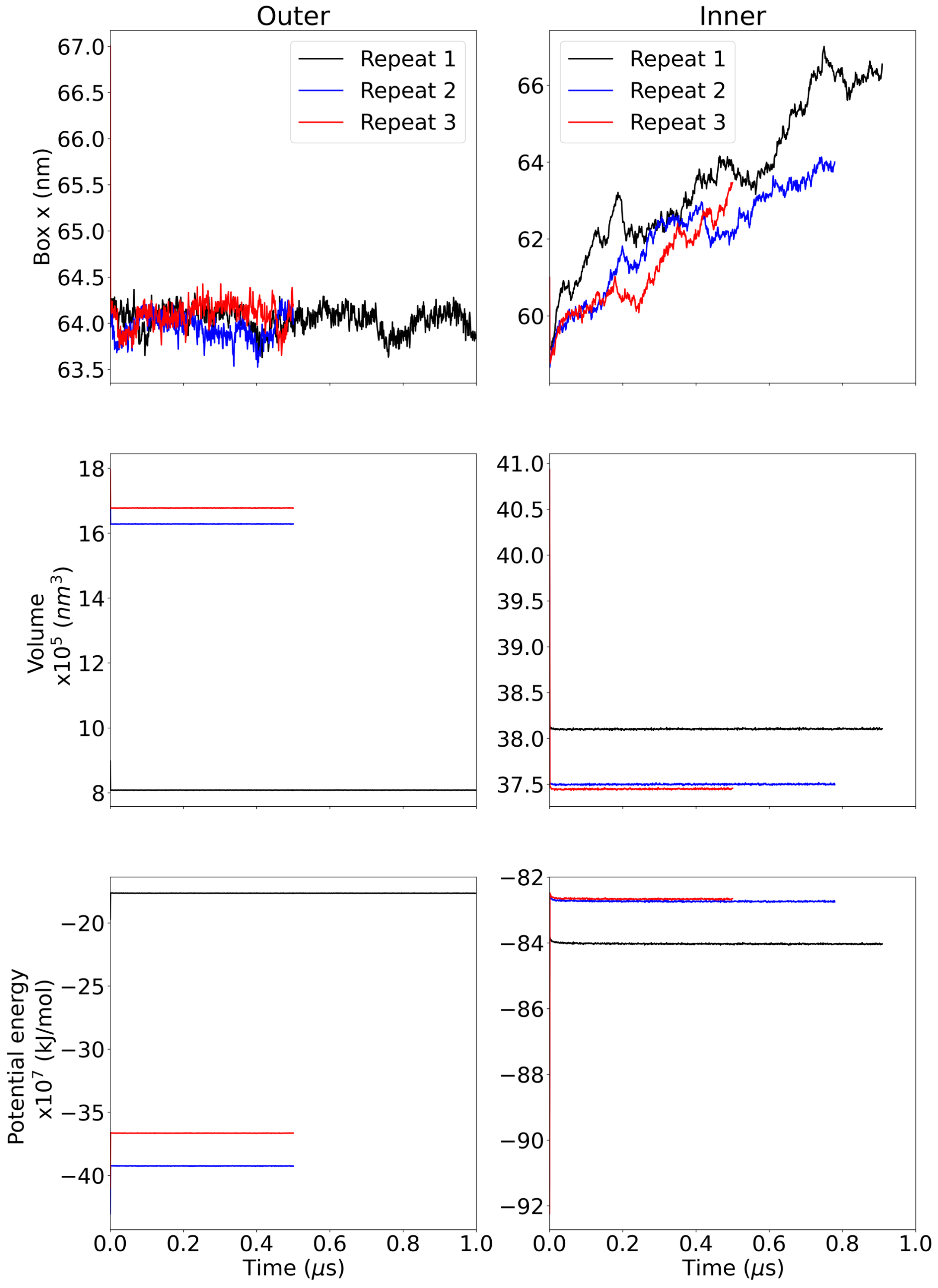


Analysis of system behaviour for membranes simulated individually. The left graphs show results from the OMM, while the right are from the IMM. Top graph shows the x box dimension, the middle graphs show the volume of the simulation box and the bottom graphs show the potential energy of the system.

**SI Figure 13**

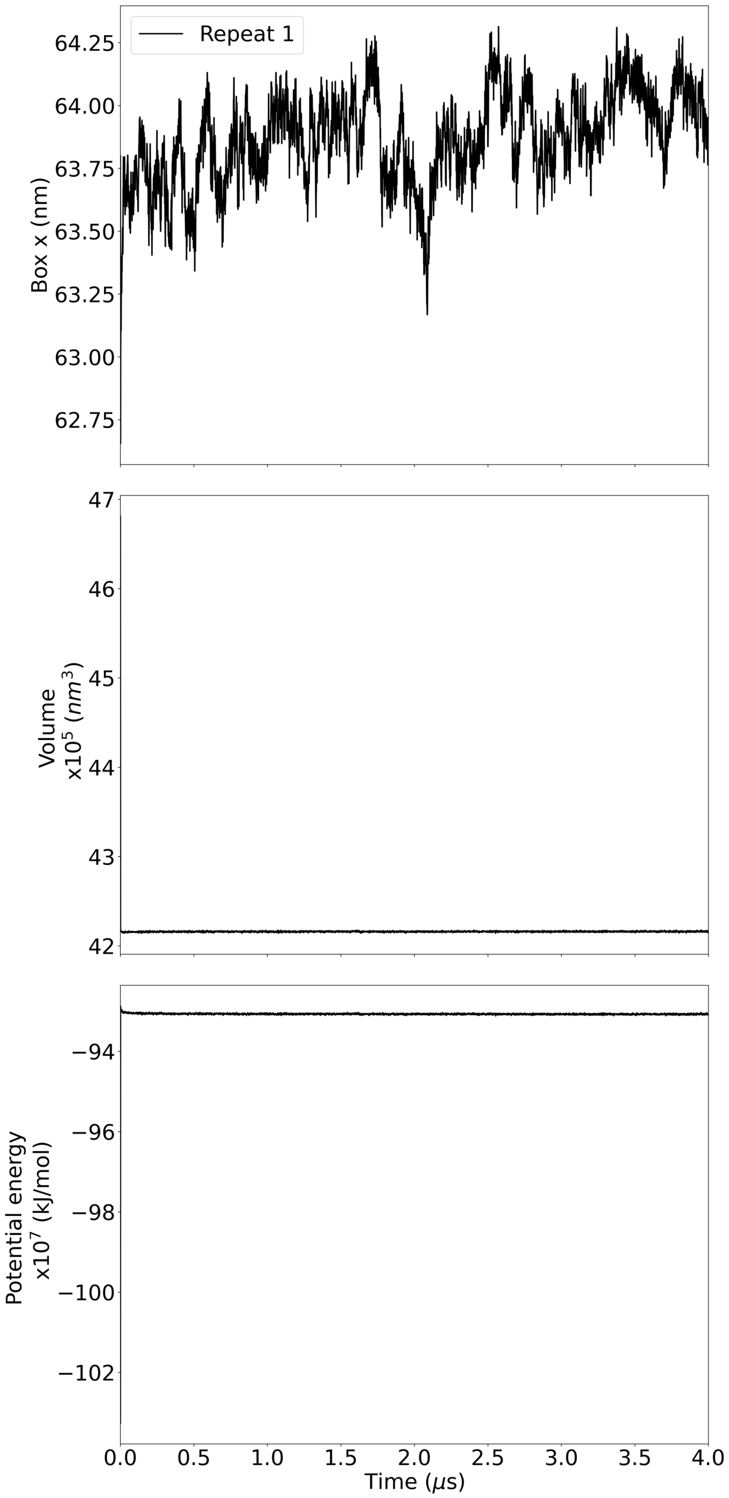


Analysis of system behaviour for the crista model. Top graph shows the x box dimension, the middle graphs show the volume of the simulation box and the bottom graphs show the potential energy of the system.
